# A cell-free platform based on nisin biosynthesis for discovering novel lanthipeptides and guiding their overproduction *in vivo*

**DOI:** 10.1101/757591

**Authors:** Ran Liu, Yuchen Zhang, Guoqing Zhai, Shuai Fu, Yao Xia, Ben Hu, Xuan Cai, Yan Zhang, Yan Li, Zixin Deng, Tiangang Liu

## Abstract

Lanthipeptides have extensive therapeutic and industrial applications; however, since many are bactericidal, traditional *in vivo* platforms are limited in their capacity to discover and mass produce novel lanthipeptides as bacterial organisms are often critical components in these systems. We developed a cell-free protein synthesis (CFPS) platform that enables rapid genome mining, screening and guiding overproduction of lanthipeptides *in vivo*. For proof-of-concept studies, the type I lanthipeptide, nisin, was selected. Four novel lanthipeptides with anti-bacterial activity were identified among all nisin analogs in the NCBI database in a single day. Further, we coupled the CFPS platform with a screening assay for gram-negative bacterial growth and identified a potent nisin mutant, M5. The titer of nisin and nisin analogs significantly improved with CFPS platform guidance. Owing to the similarities in biosynthesis, our CFPS platform is broadly applicable to other lanthipeptides, provides a universal method for lanthipeptides discovery and overproduction.

## Introduction

Lanthipeptides are a major group of ribosomally synthesized and post-translationally modified peptides (RiPPs) produced by microorganisms, characterized by intramolecular thioether crosslinks (termed lanthionine and methyllanthionine residues), with many defined biological activities (*1*). Lanthipeptides’ biosynthesis-related genes are typically assembled in gene clusters encoding a precursor peptide (LanA) that consists of a C-terminal core region and an N-terminal leader region, post-translational modification (PTM) enzymes, transporters, processing proteases, immunity proteins, and regulatory machinery (*2*). During their biosynthesis, the lanthionine and methyllanthionine residues are introduced in a two-step PTM process (*1*). In the first step, Ser and Thr residues in the core peptide of LanA are dehydrated by dehydrase. The thioether crosslinks are then formed subsequently via a Michael-type addition by Cys residues onto the dehydro amino acids. The LanA with thioether crosslinks was defined as modified precursor peptide (mLanA) and generally lack biological activities. Furthermore, following protease cleavage the leader peptide in mLanA, the remaining core peptide with thioether crosslinks, functions to exert various biological activities and is designated as mature lanthipeptides. Most lanthipeptides exhibit antimicrobial activity, hence, microbes must express transporters and immunity proteins in their biosynthetic gene cluster to achieve adequate protection.

Lanthipeptides hold much promise for discovering of novel bioactive compounds. Their biosynthetic gene clusters have been identified in the genomes of many microorganisms (*3*, *4*), however a large proportion of these microorganisms are difficult to culture in the laboratory. The heterologous synthesis of inactive mLanA (e.g., the coexpression of LanA and PTM enzymes) in *Escherichia coli* and *Lactococcus lactis* and formation of mature lanthipeptides *in vitro* provide an opportunity for lanthipeptides synthesis (*5*, *6*). Construction of a mutant lanthipeptides library and performing high-throughput screening is another useful method for discovering novel lanthipeptides. Further, method have been developed that allow for the high-throughput screening of new bioactive (anti-viral) lanthipeptides *in vivo* (*7*), and ingenious high-throughput screening of anti-microbials lanthipeptides by expression of inactive mLanAs in living cells prior to removal of the leader peptide *in vitro* (including in dead cells) to form active lanthipeptides (*8*, *9*). However, the *in vivo* systems pose unavoidable problems, such as intracellular toxicity; the formation of inclusion bodies, which is problematic for subsequent purification; lack of special tRNA^Glu^ for heterologous dehydrase (dehydration in lanthipeptide PTM process is catalyzed in a tRNA^Glu^-dependent manner). Specifically, Hudson et al. (2015) illustrated that *E. coli* tRNA^Glu^ contributes to the lack of dehydrase activity in thiomuracin (a type of RiPPs was synthesized by *Thermobispora bispora*) heterologous biosynthesis (*10*).

Besides discovering novel lanthipeptides, an urgent need exists for increased production of lanthipeptides as they have a wide range of applications in industry and medicine (*11*, *12*). It has been reported that screening for optimal strains (*13*), optimizing culture conditions (*14*), and metabolic engineering (*15*) would serve to improve the production of lanthipeptides. However, although biosynthesis of lanthipeptides occurs via the common pathway including LanA synthesis, PTM, proteolysis, and export, the rate-limiting steps of biosynthesis are unknown and no clear principle for increasing production *in vivo* has been defined.

Given that many lanthipeptides are bactericidal, cell-free protein synthesis (CFPS), which is independent of cell growth (*16*, *17*), is a promising approach for lanthipeptides research. Cheng et al. (2007) used a commercial *in vitro* rapid translation system kit to express nisin (class I lanthipeptide) precursor peptide gene (*nisA*) and PTM genes (*nisB* and *nisC*) together to form modified NisA (mNisA). The active nisin was then obtained by commercial protease trypsin treatment (*18*). Additional studies on RiPPs synthesis used cell-free systems to transcribe and translate precursor peptide genes, combined with part of original PTM purified enzymes in the RiPPs biosynthetic gene cluster and heterologous isozymes (the original one is hard to purified or could not be obtained as an active enzyme) together. One of these studies even employed a chemical reagent (H_2_O_2_ to generate dehydroalanies), to synthesize RiPPs *in vitro* (*19*, *20*). These studies demonstrate the flexibility and robustness of CFPS, however they didn’t fully reconstitute the biosynthesis of RiPPs in microbes and thus, are not capable of identifying the rate-limiting steps *in vivo* for RiPPs overproduction, or developing methods for high-throughput discovery of novel RiPPs in CFPS.

To further examine the methodological advantages of cell-free systems in the research of lanthipeptides, we developed a CFPS platform that address the current issues faced in genome mining, screening of novel antimicrobial lanthipeptides, and achieving overproduction of lanthipeptides via CFPS platform guidance (Fig. 1). Nisin was selected for our proof-of-concept experiments. It is the first commercially available lanthipeptide, classified as type I, that has been used as a food preservative worldwide for more than 60 years without the development of bacterial resistance, and has a well-defined catalytic mechanism (*6*, *21*, *22*). We developed an *E. coli* CFPS platform using a simple preparation method to fully reconstruct the nisin biosynthetic pathway for *in vivo* biosynthesis and optimization for higher efficiency. To test the functionality of this CFPS platform, we first performed genome mining on all potential nisin analogs in the NCBI database using our CFPS platform in a single day. We next developed a screening process for the identification of lanthipeptides that are functionally active against gram-negative bacteria and applied it to our CFPS platform to assess the nisin mutant library. Thirdly, we used one-step metabolic engineering *in vivo* to overproduce nisin in an industrial host and nisin analogs in a heterologous expression host; the engineered target was identified using the optimized nisin CFPS platform.

**Fig. 1.**
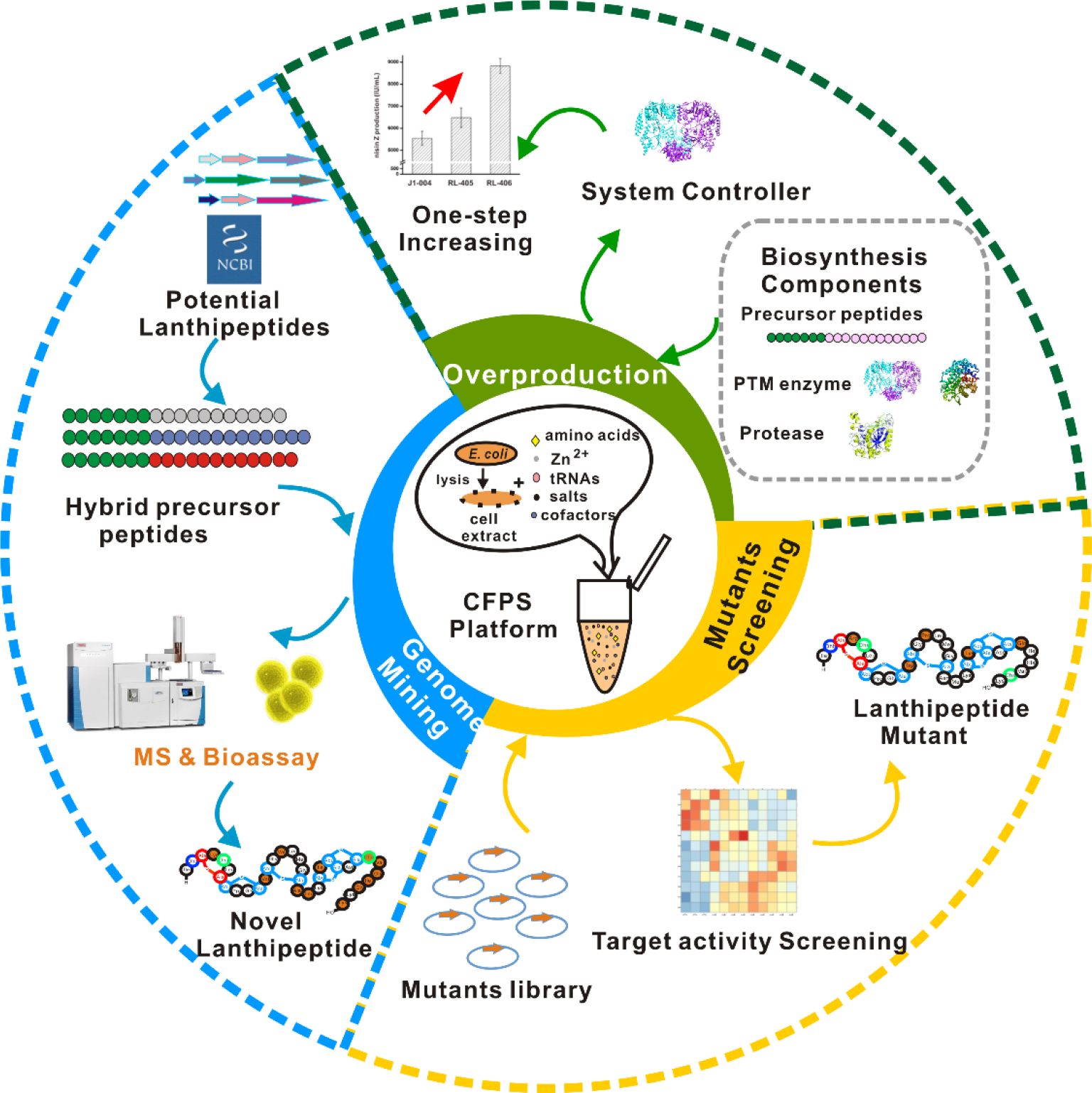
CFPS platform for genome mining, mutants screening, and guiding *in vivo* overproduction of lanthipeptides. The blue dotted area represents genome mining of potential lanthipeptides using the CFPS platform by a hybrid precursor peptide strategy. The yellow dotted area represents the screening of lanthipeptides mutants by target activity screening using the CFPS platform. The green dotted area represents the *in vitro* reconstruction of the biosynthetic pathway of nisin using the CFPS platform and identifies key system controllers to guide the overproduction of nisin *in vivo*.

## Results

### A cell-free protein synthesis platform for lanthipeptide biosynthesis

Although commercial nisin, including nisin A and nisin Z, contain differences in their structure namely at the amino acid residue in position 27, no significant differences have been described in their functions. The modified nisin A precursor peptide (mNisA) has been successfully synthesized by a commercial *E. coli*-based CFPS, and the active nisin was obtained by commercial protease trypsin cleavage of the leader peptide in mNisA (*18*). However, since the biosynthetic system of nisin is not completely reconstituted in CFPS (lacks original protease NisP), it cannot be used to identify the rate-limiting step of nisin biosynthesis *in vivo*, nor can it be used to enhance the CFPS efficiency for nisin synthesis by systematic titration of nisin’s components (precursor peptide NisA, PTM enzyme NisB and NisC, and protease NisP), while using commercial kits to perform large-scale screening is costly. Therefore, we prepared a basic form of the CFPS system in laboratory (*23*) for the expression of nisin biosynthetic enzymes from a DNA template. This CFPS system included *E. coli* cell extracts lacking any remaining living cells (Fig. S1) and 26.5 mg/mL lysate protein supplemented with Zinc ions to facilitate the activation of cyclase, which is responsible for the thioether crosslink formation. Four plasmids were constructed (pJL1-*nisZ* for expression of precursor peptide NisZ; pET28a-*nisB* and pET28a-*nisC* for expression of PTM enzymes NisB and NisC, respectively; and pET28a-*nisP* for expression of protease NisP) to achieve nisin biosynthesis in our CFPS system.

We evaluated the performance of our CFPS platform for nisin biosynthesis. Each plasmid (0.5 nM) (pJL1-*nisZ*, pET28a-*nisB*, pET28a-*nisC*, and pET28a-*nisP*) was added into the CFPS platform and incubated for 6 h (Fig. 2A). The 400 μL reaction mixture was concentrated to 20 μL and the concentrated mixture was then detected by tandem mass spectrometry (LC-MS-MS). The results confirmed that fully modified nisin Z was produced in the CFPS mixture (Fig. 2B). The concentrated mixture was then used for antibacterial bioassays using the agar diffusion method, which revealed an obvious zone of inhibition on a *Micrococcus luteus* plate (Fig. S2A). This result confirmed that nisin Z synthesized by the CFPS platform was biologically active.

**Fig. 2.**
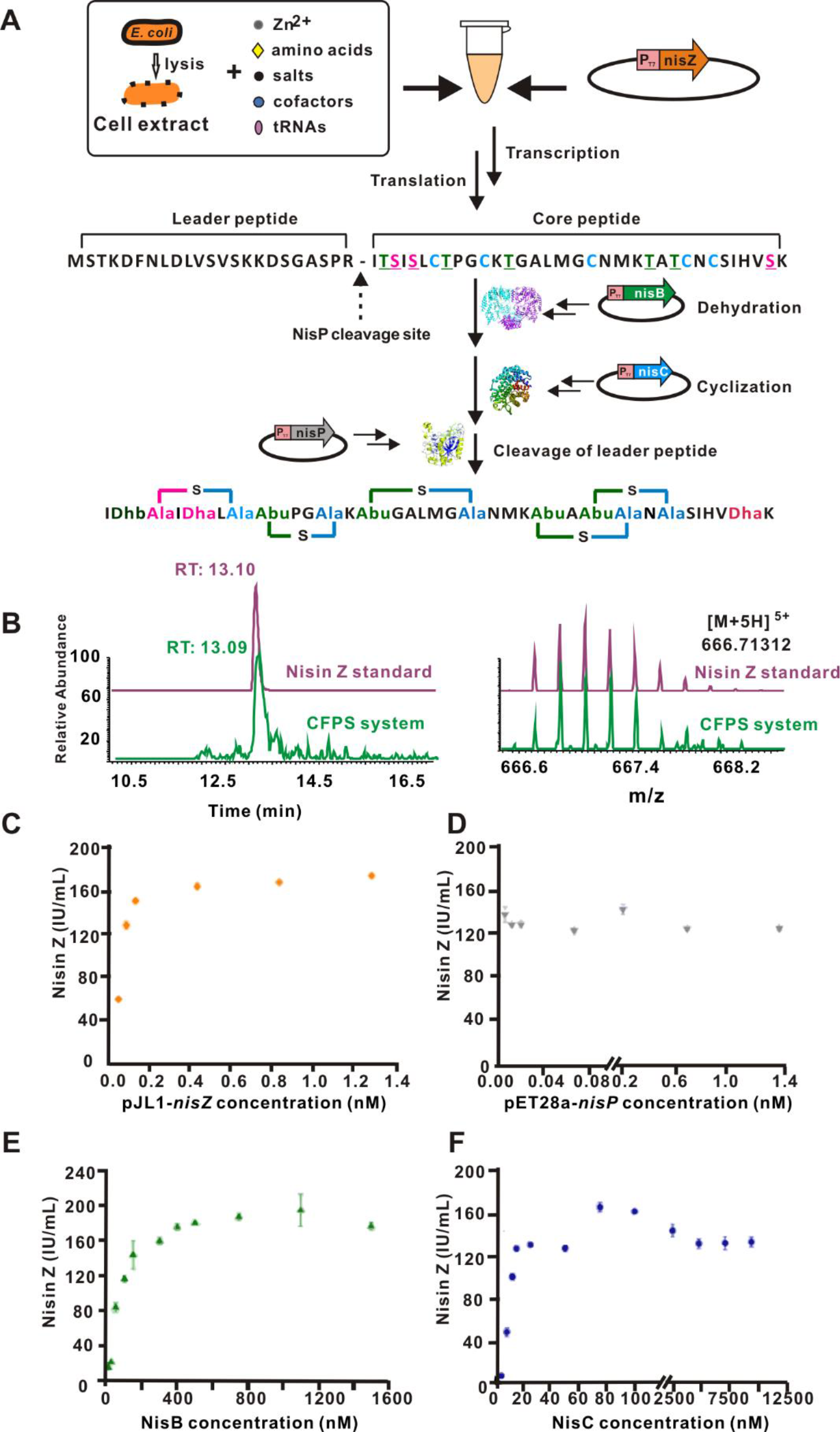
Reconstitution and optimization of nisin biosynthesis using the CFPS platform. **(A)** Schematic illustration of reconstitution of nisin Z in CFPS (Dha: dehydrolalanine; Dhb: dehydrobutyrine; Abu: 2-aminobutyric acid). (**B**) LC-MS-MS qualitative assay of nisin Z CFPS and standard, [M+5H]^5+^ ion of nisin Z (m/z = 666.71312) was used for ion monitoring. (**C**) Titration of pJL1-*nisZ* involved in the nisin biosynthesis machinery. **(D)** Titration of pET28a-*nisP* involved in the nisin biosynthesis machinery. **(E)** Titration of NisB involved in the nisin biosynthesis machinery. **(F)** Titration of NisC involved in the nisin biosynthesis machinery. Error bars are based on three independent replicates.

### Optimization of the nisin CFPS platform

Although bioactive nisin was synthesized using the initial CFPS platform, the efficiency was low. Therefore, to enhance the CFPS efficiency of nisin, we used western blotting to analyze the accumulation of nisin biosynthetic enzymes and identified the system controller of the initial nisin CFPS platform. We detected a single hybridization band at the expected size of the precursor peptide (Fig. S2B), indicating that the other three enzymes responsible for nisin PTM (NisB and NisC) and leader peptide cleavage (NisP) were poorly expressed. Accordingly, his6-tagged NisB, NisC, and NisPs (truncated NisP) (*22*) were overexpressed in *E. coli* and purified. The system controller was then investigated by systematically replacing each plasmid with 500 nM of the corresponding purified protein. The substitutions of NisB and NisC significantly increased production of nisin Z (quantitative unit IU, 40 IU=1 μg nisin (*24*)); while the replacement of pJL1-*nisP* with NisPs had minimal effects on the production of nisin (Fig. S2C). These results indicate that the dehydration and cyclization of the precursor peptide, controlled by NisB and NisC, serves as the system controller in the nisin PTM process. Moreover, the cleavage of the leader peptide did not act as a bottleneck.

We used systematic titration to study the optimal concentration of each component in the nisin biosynthetic pathway using the CFPS platform. First, concentrations of the precursor peptide gene encoded by plasmid pJL1-*nisZ* were examined by varying the concentration, while fixing the concentrations of NisB and NisC to 0.5 μM and pET28a-*nisP* to 0.5 nM. The highest level of nisin Z production from pJL1-*nisZ* was detected for concentrations of 0.4–1.3 nM (Fig. 2C). A similar titration for pET28a-*nisP* was applied to examine the effect of *nisP* concentration on the CFPS platform. Consistent with the results of our previous enzyme replacement experiments, the concentration of pET28a-*nisP* did not appreciably influence nisin production (Fig. 2D). Based on the *nisZ* and *nisP* titration studies, the optimal concentrations were determined to be 1.3 nM and 0.1 nM, respectively.

We next investigated the two enzymes responsible for nisin-induced PTMs. In these assays, concentrations of pJL1-*nisZ* and pET28-*nisP* were set to 1.3 nM and 0.1 nM, respectively. When testing the dehydration reaction, NisC was set to 500 nM, and the concentration of NisB was varied from 10 to 1,500 nM. The production of active nisin Z improved dramatically (from ~15 IU/mL to ~180 IU/mL) when the NisB concentration increased from 10 nM to 800 nM (saturation occurred at concentrations above 1,000 nM NisB; Fig. 2E). Thus, increasing the concentration of NisB over a larger range would help increase nisin production. Moreover, a high NisB concentration did not inhibit nisin Z production. These results confirmed that overexpression of *nisB* contributed substantially to nisin overproduction. When the NisB concentration was 500 nM, the production of active nisin Z was observed from ~10 IU/mL to ~165 IU/mL when NisC was varied from 1 nM to 80 nM; a higher NisC concentration (from 100 to 10,000 nM) did not appreciably influence nisin Z production (Fig. 2F). Overall, the nisin titer of CFPS increased by more than 30-fold (from <5 IU/mL to ~180 IU/mL) following replacement of nisin PTM enzyme, and the optimized CFPS platform was used in subsequent studies.

### Use of the CFPS platform for rapid genome-mining of novel lanthipeptides

In a previous genome mining study, the core peptide of a potential lanthipeptide was fused to the nisin leader peptide to form a hybrid precursor peptide, which were then co-expressed with *nisBC* and *nisT* (transporter) in *Lactococcus lactis*. The leader peptides of mLanAs were removed by NisP or trypsin *in vitro* to form bioactive lanthipeptides. Five novel lanthipeptides similar to nisin with anti-bacterial activity were identified (*6*). Here, we explored the use of our CFPS platform for mining potential in lanthipeptides (nisin analogs) with anti-bacterial activity using the same nisin modification machinery.

We designed a rapid genome mining scheme (Fig. 3A). All proteins with 40-80 amino acids in length were selected from the NCBI database (accessed June 2018). The resulting number of proteins was over two million. The sequence “SxSLCTPGCxTG” (where x denotes an arbitrary residue) was used to retrieve all potential nisin analogs. A total of 210 analogs were identified. We then excluded known lanthipeptides by alignment to sequences in the BAGLE4 database (*25*) and found 18 potential lanthipeptides (RL1–RL18; Table 1). Their core peptides were linked to the nisin leader peptide by gene synthesis to form hybrid precursor peptides (Table S5).

**Fig. 3.**
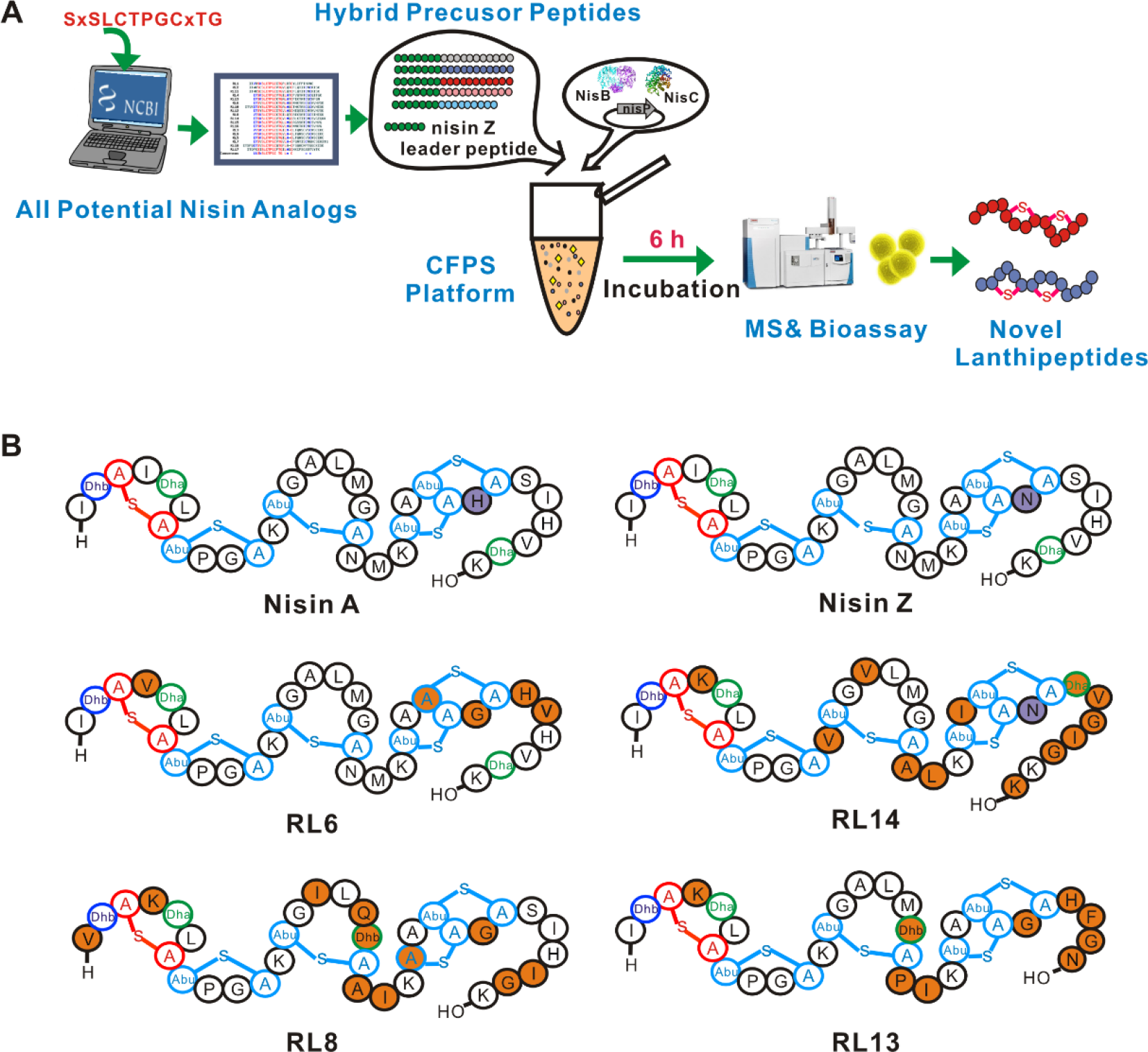
Rapid genome mining for novel lanthipeptides in CFPS. **(A)** Schematic illustration of rapid genome mining for novel lanthipeptides using the CFPS system. **(B)** Nisin and novel nisin analogs. Dha: dehydrolalanine; Dhb: dehydrobutyrine; Abu: 2-aminobutyric acid

**Table 1.**
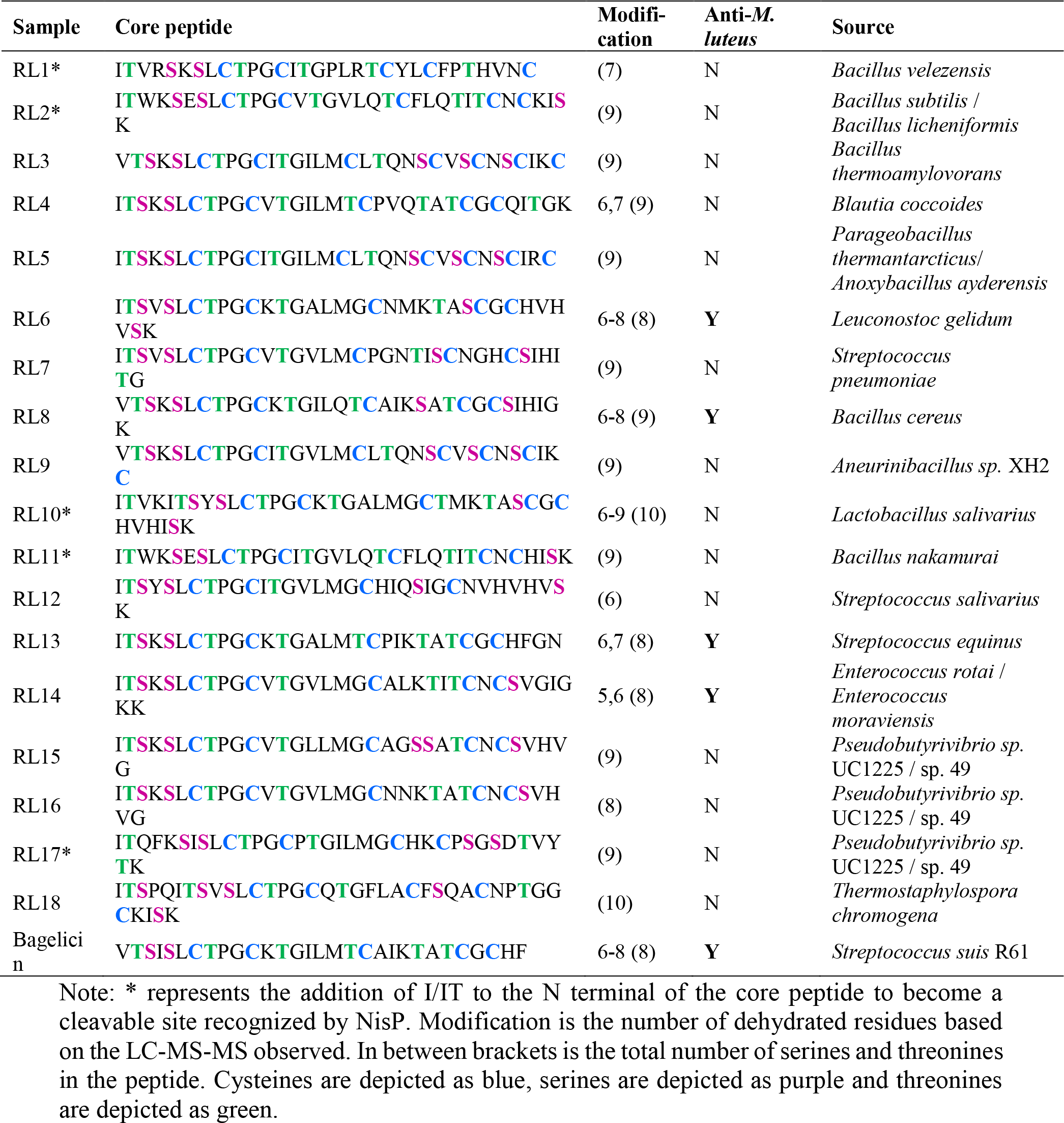
List of identified nisin analog candidates.

These hybrid precursor peptides were then cloned into the pJL1 plasmid and bagelicin, a screened nisin analog with anti-bacterial activity (*6*), was chosen as the positive control for validation. The pJL1-*nisZ* plasmid in our CFPS platform was replaced with the hybrid precursor peptide plasmids (pRL1–pRL18, pJL1-bagelicin). After incubation for 6 h, the bagelicin CFPS mixture (positive control) was analyzed by LC-MS-MS and bioassays. The results indicate that the Thr and Ser residues in bagelicin were dehydrated and the bagelicin CFPS mixture contained anti-*M. luteus* activity (Table 1), demonstrating the successful application of our CFPS platform to genome mining, as previously reported using *in vivo* methods (*6*). Six candidate lanthipeptides were dehydrated based on LC-MS-MS detection, among which four displayed anti-*M. luteus* activity (Table 1).

To characterize the structure and antibacterial ability of novel lanthipeptides, we prepared mLanAs from novel lanthipeptides (RL6, RL8, RL13, and RL14) in *E. coli* (Table S1) and removed their leader peptides *in vitro*. Since the structure of nisin was determined using its most effective antibacterial component (eight-fold dehydrated components) (Fig. S3), the eight-fold dehydrated component of RL6, RL8, RL13, and RL14 were also used to determine their structure (Fig. 3B, Fig. S4). We treated the novel lanthipeptides with thiol-alkylating reagents *N*-ethylmaleimide (NEM) to sequester any noncyclic Cys residues and evaluated ring topology by LC-MS (Fig. S5), the results showed all five thioester rings in their core peptides were formed in mature lanthipeptides. We then tested the antibacterial activity of novel lanthipeptides, including activity against *M. luteus* and clinical pathogenic strains of *Enterococcus faecalis*, *Staphylococcus aureus*, and methicillin-resistant *Staphylococcus aureus* (MRSA). All four tested lanthipeptides exhibited antibacterial activity against these bacteria, with RL14 outperforming nisin against *M. luteus* and *E. faecalis* based on the quantification of lanthipeptides using the eight-fold dehydrated molecules (Table S2).

### Screening of lanthipeptides activity using the CFPS platform

Next, we evaluated the performance of the CFPS platform for library screening, an application that has been reported using *in vivo* methods (*7*–*9*). We screened mutant lanthipeptides using the CFPS platform to extend the specific activity of nisin against gram-positive bacteria to gram-negative bacteria. The screening of mutant nisin with anti-gram-negative bacteria has been reported by the knockout of *nisA* in a nisin A producing strain of *L. lactis* and the introduction of a plasmid harboring *nisA* mutants. Several nisin mutants (including S29A) with anti-*E. coli* activity were identified using this approach (*26*).

We aligned reported nisin analogues (nisin A, nisin Z, nisin F (*27*), nisin Q (*28*), and nisin U (*29*)) and selected five amino acid residues in non-conserved regions (positions 4, 12, 15, 24, and 29 of nisin Z) for saturation mutagenesis to form a mutant nisin library. The nisin mutants were synthesized using our optimized CFPS platform on microplates with 6-h incubation periods. The CFPS reaction mixture was then co-cultured with *E. coli* in 96-well plates, and the OD_600_ of the culture was determined after 8 h. The mutant with lowest OD_600_ inhibited the growth of *E. coli* (Fig. 4A). In order to improve the accuracy of the screening, we conducted two rounds of screening. In the first round, the candidates that effectively inhibited the growth of *E. coli* were selected and compared with nisin in the second round of screening. The mutants that elicited more significant levels of *E. coli* growth inhibition than nisin were selected. After two rounds of screening, 2 out of 3,000 mutants showed stronger inhibitory effects against *E. coli* than did nisin (Fig. 4C). By sequencing the two plasmids, the M4 mutant was identified and named based on the four mutations in its core peptide (I4K, P9T, K12L, and S29D). Additionally, the M5 mutant was identified and named based on the five mutations in its core peptide (I4R, K12W, A15P, A24K, and S29Q).

**Fig. 4.**
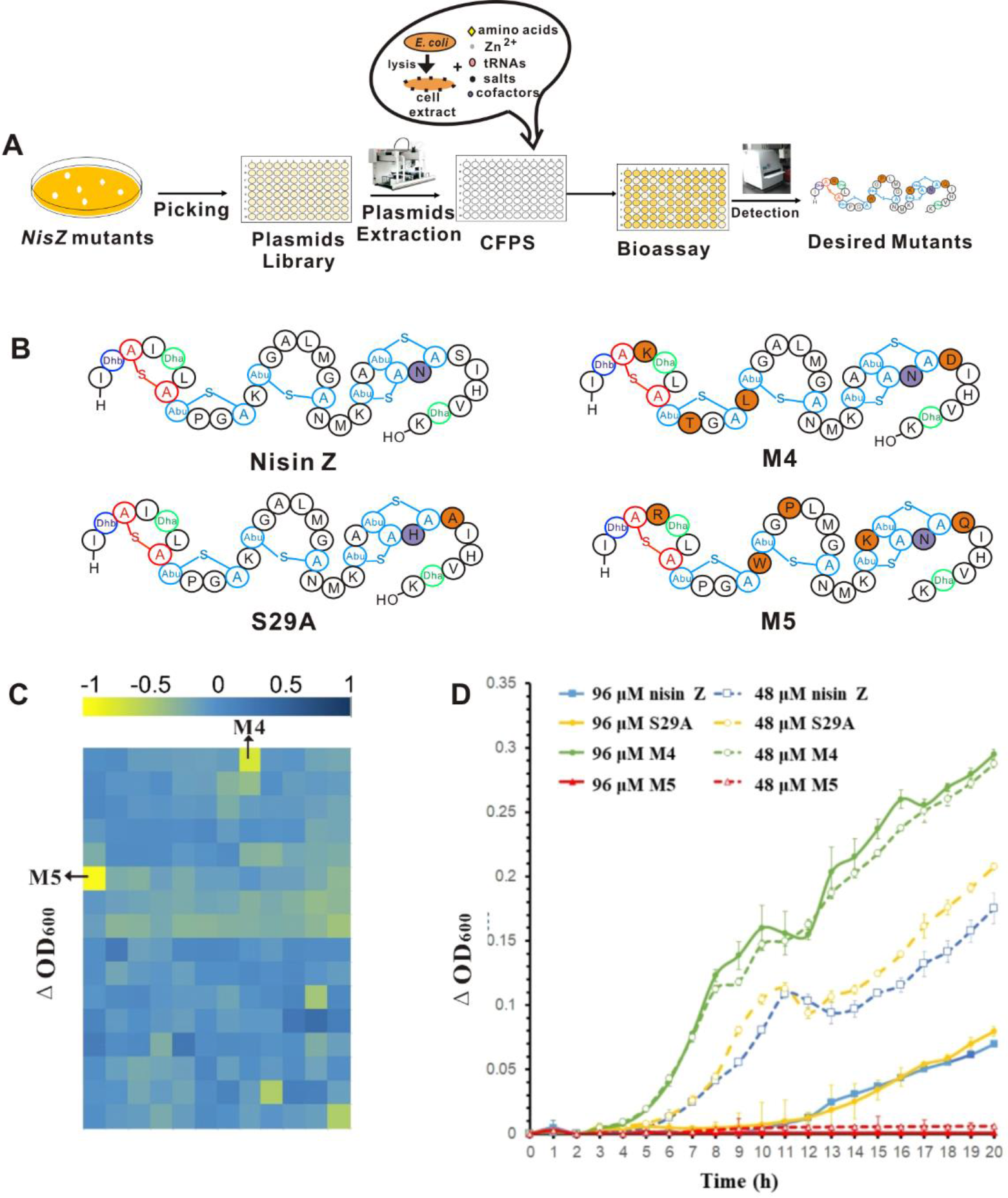
Screening of nisin mutants using the CFPS system. **(A)** Schematic illustration of the screening of nisin mutants. **(B)** Nisin and nisin mutants. Dha: dehydrolalanine; Dhb: dehydrobutyrine; Abu: 2-aminobutyric acid (**C**) Heat-map of antimicrobial activity of nisin mutants against *E. coli*. ΔOD_600_ values of *E. coli* for the nisin mutant subtracted from the OD_600_ value for the control group (nisin action) in the same 96-well plate are shown. Yellow dot represents an inhibitory effect on *E. coli*. (**D**) OD_600_ of *E. coli* DH5α in LB medium with 320 μM EDTA after treatment with nisin and nisin mutants. ΔOD_600_ is the difference between readings with different concentrations of blank media. Error bars are based on three independent replicates.

Next, to verify the reliability of the CFPS platform for screening, we purified M4, M5, and S29A using the method for the expression of mLanAs in *E. coli* (Table S1) and removed the leader peptide *in vitro*. The structures were identified by LC-MS-MS (Fig. 4B; Fig. S4) and the thioester rings were detected by NEM reaction (Fig. S5). To ensure similar conditions to those used in previous reports so that nisin exhibited observable inhibition against *E. coli* (*26*, *30*, *31*), DH5α with 320 μM EDTA was used based on the apparent growth inhibition of *E. coli* with different concentrations of nisin (Fig. S6). We generated the *E. coli* DH5α growth curve using M4, M5, S29A, or nisin Z and found that M5 inhibited growth more effectively against gram-negative *E. coli* than did nisin Z or S29A (Fig. 4D). These results suggest that our novel CFPS platform is effective in screening for lanthipeptides.

### Targeted metabolic engineering for nisin and nisin analogs overproduction in vivo

We have learned from the process of optimizing the CFPS platform that increasing NisB significantly increased nisin Z production. We next evaluated this strategy using industrial *L. lactis* strains for nisin Z overproduction. The industrial nisin Z producing strain is difficult to manipulate genetically since it was mutated for decades, making its ability for accepting foreign DNA poor. Hence, the specific genetic operation employed is critical when working with this industrial strain. To verify whether the overexpression of *nisB in vivo* could increase nisin production, RL405 for overexpression of *nisZ* and RL406 for co-overexpression of *nisZ* and *nisB* were constructed (Fig. 5A). The results (Fig. 5C) show the industrial strain J1-004 produced 5,549.0 ±316.3 IU/mL of nisin Z after 16 h of fed-batch fermentation, while RL405 produced 6,479.7 ±443.9 IU/mL of nisin Z under the same fermentation conditions, which was 16.8% higher than that of J1-004. However, the *nisB* overexpressed strain, RL406, exhibited an increase in nisin Z production to 8,828 ±336.2 IU/mL, representing a nearly 60% increase over that of the industrial strain J1-004 and a 36.2% improvement to RL405. This result further demonstrated that the overexpression of *nisB* contributed substantially to nisin overproduction. We further analyzed the nisin biosynthesis proteins in the engineered *L. lactis* strains by a targeted proteomics approach. NisZ was detected in RL405 (both NisZ and NisB were detected in RL406) and *nisZ* and/or *nisB* were overexpressed, as designed (Fig. S7).

**Fig. 5.**
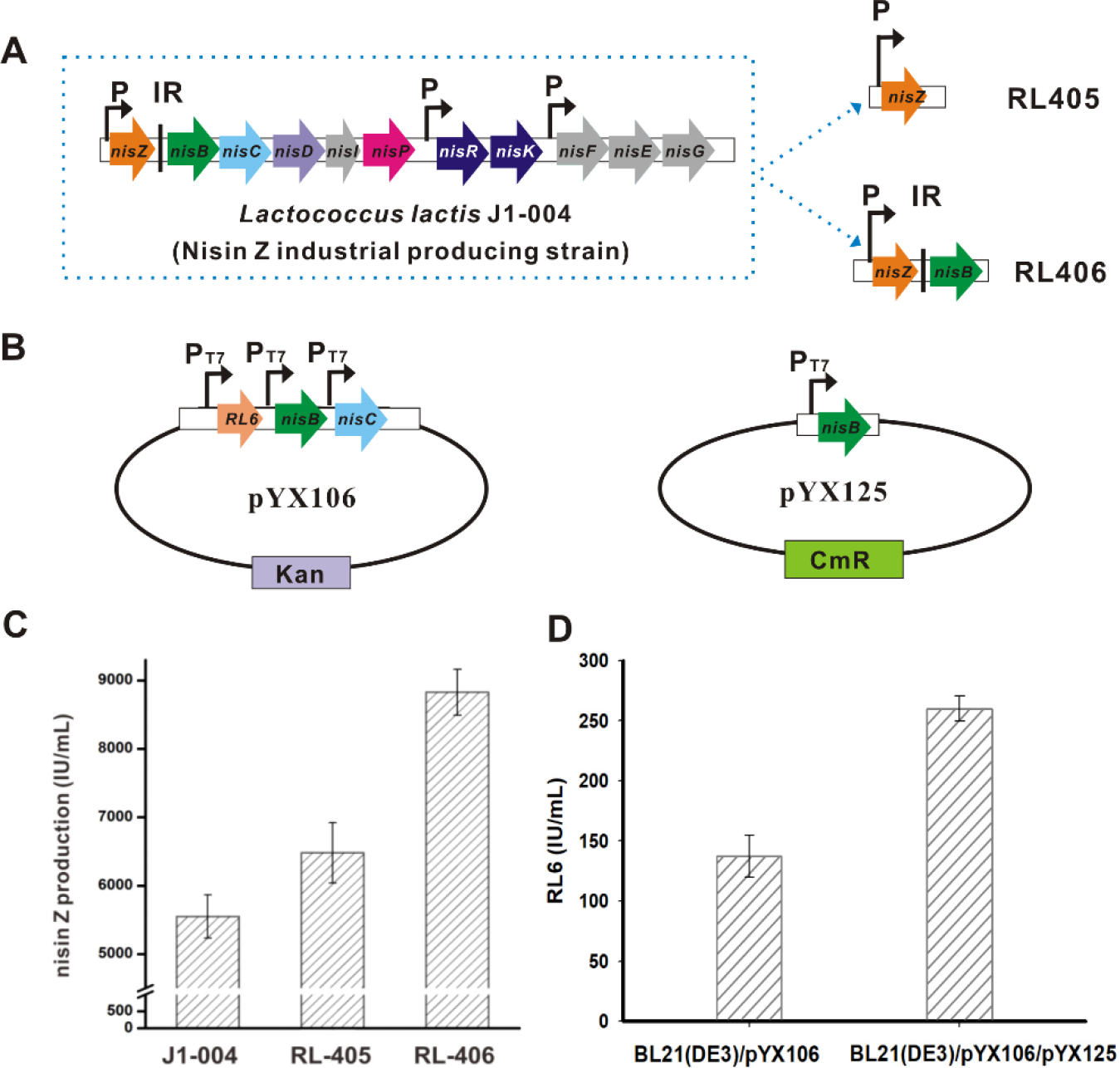
CFPS platform guides the overproduction of lanthipeptides. **(A)** Original nisin biosynthesis gene cluster in *L. lactis* J1-004 and construction of the overexpression operons used in this study. (**B**) Plasmids for the expression of the modified RL6 precursor peptide in *E. coli*. (**C**) Production of nisin in industrial strains and the individual engineered strain after 16 h of fed-batch fermentation. (**D**) Novel lanthipeptide RL6 production in different *E. coli* strains. These two *E. coli* strains were fermented to overexpress modified RL6 precursor peptides and digested with trypsin *in vitro*. Nisin was used as the quantitative standard. Error bars are based on three independent replicates.

To investigate whether the overexpression of *nisB* could also increase the yield of nisin analogs in a heterologous expression host, *E. coli*, BL21(DE3)/pYX106 (Table S1) was constructed according to a previously described method (*5*) with *lanA* of RL6, *nisB*, and *nisC* co-expression in pYX106 for the production of mLanA of RL6 (mRL6) (Fig. 5B). BL21(DE3)/pYX106/pYX125 (Table S1) was constructed for the overproduction of mRL6 with an extra copy of *nisB* expression in pYX125 besides pYX106 (Fig. 5B). The parallel fermentation of these two strains resulted in no obvious differences in OD_600_ values, indicating that the overexpression of *nisB* had no effect on the growth of *E. coli*. After cleavage of the leader peptide *in vitro* by trypsin, the *nisB* overproduction strain BL21(DE3)/pYX106/pYX125 produced RL6 at 260.1 IU/mL, while the control strain BL21(DE3)/pYX106 produced 137.5 IU/mL. The overexpression of *nisB* increased the titer by 89.2% in *E. coli* (Fig. 5D), confirming *nisB* is a universal key determinant for nisin analogous lanthipeptides overproduction. We can conclude from these results that the CFPS platform guided successful overproduction of lanthipeptides.

## Discussion

We developed an optimized CFPS platform for the genome mining of nisin analogs, screening of mutants, and guiding lanthipeptides overproduction *in vivo*. The functionality of this platform was verified in that all nisin analogs with bactericidal effects were mined using our CFPS platform in a single day, one of which (RL14) exhibited stronger anti-bacterial activity than that of nisin. Moreover, a nisin mutant (M5) with activity against gram-negative bacteria was screened using this platform; and a 60% nisin Z increase in an industrial host and an 89.2% increase in nisin analogs in a heterologous expression host.

The first advantage of our CFPS platform for lanthipeptide research is efficiency. Although we used purified nisin PTM enzymes protein in the CFPS platform for improved efficiency, these enzymes can be purified in abundance and stored at −80 °C for an extended period of time, allowing lanthipeptides to be synthesized from DNA templates in 6 h. The second advantage of our CFPS platform is that the production of lanthipeptides is not dependent on cell growth. To ensure that it has the advantage of screening for bacteriostatic activity of lanthipeptides, we developed the process of screening lanthipeptide with anti-*E. coli* activity. However, by changing the indicator strain, the platform can be used for screening other bacteriostatic lanthipeptides as well. The third advantage of our CFPS platform is that the reaction volume is 8 μL, and the reaction mixture can be processed for use in bioassays and MS detection (after desalting) without the requirement for complex purification processes. It is, therefore, reasonable to suggest that the CFPS platform can perform high-throughput screening work. The fourth advantage of our CFPS platform is the rapid and accurate identification of the system controller, allowing for identification of complex regulatory mechanisms in the cell so that the rate-limiting steps of biosynthesis can be more intuitively identified.

We provide an effective method for the cell-free production of lanthipeptides, with applications for the biosynthesis of other RiPPs. The direct synthesis of lanthipeptides using DNA templates is less efficient than synthesis using purified enzymes; however, the development of cell-free systems with higher yields (*32*, *33*), lower costs (*34*), and simpler preparation methods (*35*) is an active area of research. These studies will contribute to increasing the efficiency of directly synthesized RiPPs using DNA templates, and allow CFPS to play a more prominent role in RiPPs research.

In the CFPS platform, we used PTM enzymes (NisB and NisC) with clear catalyzed machinery to modify other potential lanthipeptides (using hybrid precursor peptides). Although novel lanthipeptides may not share the same natural structure as their original structure *in vivo*, this method represents a general and rapid approach for genome mining for potential RiPPs. More importantly, owing to the diversity of PTMs, different types of PTM enzymes can be applied to generate novel RiPPs in a combinatorial biosynthesis manner (*36*, *37*). Due to the robustness and flexibility of CFPS, different PTM enzymes can readily be combined in this cell-free system to achieve novel RiPPs with diverse bioactivities.

CFPS can be easily developed as an automated high-throughput screening platform. The Freemont group at ICL has developed a rapid automated method using the cell-free platform to quantify a series of ribosome-binding site mutants and uncharacterized endogenous constitutive and inducible promoters to characterize new DNA components in non-model bacteria (*38*). It is, therefore, reasonable to suggest that combined automated high-throughput platforms containing CFPS with different microbes, such as *Streptomyces* and cyanobacteria which possess many RiPP gene clusters, will accelerate the rate of discovery for novel RiPPs since their cell extracts may contain novel components related to the synthesis of RiPPs. Overall, our research extends the use of cell-free systems to address the issue facing lanthipeptides overproduction and novel compound discovery, while providing the possibility for development of CFPS platforms for other RiPPs studies.

## Materials and Methods

### Strains

The strains used in this study are listed in Table S1. *E. coli* was grown in LB with appropriate antibiotics (Kan: 50 μg/mL, CmR: 34 μg/mL) at 37°C. *L. lactis* was grown in M-17 (Oxoid, Thermo Fisher Scientific, Waltham, MA, USA) supplemented with 0.5% D-glucose (GM-17) at 30°C with appropriate antibiotics (EmR: 5 μg/mL). *M. luteus* was grown in bioassay medium with 1.2% tryptone, 0.75% yeast extract, 0.75% NaCl, 0.3% NaH_2_PO_4_, and 0.75% D-glucose at 37°C. *E. faecalis*, *S. aureus*, and MRSA were grown in Mueller–Hinton broth (MHB) (BD Difco, Franklin Lakes, NJ, USA) at 37°C.

### Plasmids

The primers and the plasmids for strain construction are listed in Tables S3 and S4, respectively. In general, the primers used to construct the relevant plasmids were named after the plasmid, and the corresponding restriction enzyme sites were designed on the primers. For instance, pJL1-nisZ-F and pJL1-nisZ-R were used for *nisZ* amplification and the PCR fragment cloned into the *Nde*I and *Bam*HI sites of pJL1 (*39*) to yield pJL1-*nisZ*. All fragments obtained by polymerase chain reaction were gel-purified using a DNA Gel Extraction kit (Axygen, Corning, NY, USA) according to manufacturer’s instructions, before cloning.

For the biosynthesis of nisin and other lanthipeptides in the CFPS system, genomic DNA from *L. lactis* was obtained using the Blood & Cell Culture DNA Mini kit (QIAGEN, Hilden, Germany) following the manufacturer’s instructions. *nisZ* was amplified by PCR from *L. lactic* J1-004, subcloned to pJL1 (*39*), and named pJL1-*nisZ*. *nisB*, *nisC, nisP*, and *nisPs* were individually amplified by PCR from *L. lactic* J1-004 and cloned to pET28a(+) (Novagen, Darmstadt, Germany) to yield pET28a-*nisB*, pET28a-*nisC*, pET28a-*nisP*, and pET28a-*nisPs*, respectively. Each hybrid lanthipeptide RL1–RL18 (see Table S5 for sequences) was amplified using pRL-F as the general forward primer and pRLX-R as the reverse primer (X represents a specific plasmid; Table S3). For example, the reverse primer for pRL1 is pRL1-R, followed by the PCR fragment cloned into the *Nde*I and *Bam*HI sites of pJL1-*nisZ*, respectively, yielding pRL1–pRL18.

For the expression of mLanAs in *E. coli*, several plasmids were constructed. An ~3 kb *Nde*I/*Kpn*I fragment containing *nisB* was inserted into the corresponding sites of the plasmid pRSFDuet-1 and yielded plasmid pYZ82. In the plasmid pYZ85, the *lanA* of RL6 was inserted between the *Bam*HI and *Eco*RI sites of the plasmid pYZ82, which was constructed to express the *lanA* (RL6) gene with a His6-tag and *nisB*. Similarly, pYZ86, pYZ87, pYZ89, pYZ90, and pYZ91 were constructed by inserting the *lanA* (RL8, S29A, M5, M4, and NisZ) genes between the *Bam*HI and *Eco*RI sites of the plasmid pYZ82, respectively. Five plasmids, pYZ92, pYZ93, pYZ95, pYZ96, pYZ97, and pYZ99, were constructed to replace the His6-tag in pYZ90, pYZ89, pYZ85, pYZ86, pYZ87, and pYZ91 with the sumo-tag. Primers Sumo-F and Sumo-R were used to amplify the sumo-tag fragment from pSUMO. The plasmid pYZ81 for the expression of *nisC* under the control of T7 promoter used an ~1.25 kb *Kpn*I/*Xho*I fragment inserted into the corresponding sites of the plasmid pACYCDuet.

For the expression of the mLanAs in *E. coli* with *nisB* overexpression, several plasmids were constructed. The 1.5-kb *Xho*I/*Xho*I fragment that contained the T7 promoter and *nisC* was inserted into the corresponding sites of the plasmid pYZ85, yielding pYX106. The plasmid pYX125 for the expression of *nisB* under the control of the T7 promoter used an ~3 kb *Kpn*I/*Xho*I fragment inserted into the corresponding sites of the plasmid pACYCDuet. The plasmid pYX105 was constructed to produce a sumo-tag S29A precursor peptide. An ~0.2 kb *Bam*HI/*Eco*RI fragment of a different *lanA* was inserted into the corresponding sites of the plasmid pYX105, yielding pYX122 (sumo-tag-mRL13) and pYX123(sumo-tag-mRL14). For the plasmid pYX126, *nisB* under the control of the T7 promoter was inserted into *Xho*I site of the plasmid pYZ81 for the expression of *nisB* and *nisC*.

For overexpression of lanthipeptides in *L. lactis*, two plasmids were constructed via the Gibson assembling method (*40*). The plasmid pRL415 overexpressing *nisZ* under the control of Pnis promoters was constructed. Primers pRL415-F and pRL415-R were used for the amplification of the Pnis-*nisZ* operon from *L. lactic* J1-004. Primers pRL415-VF and pRL415-VR were used for the amplification of the pMG36e backbone. The plasmid pRL423 for the overexpression of *nisZ* and *nisB* under the control of the Pnis promoter was constructed by co-amplification in their original order in the *nisZ*BTCIP operon (*41*). Primers pRL423-F and pRL423-R were used for the amplification of *nisB* from *L. lactic* J1-004. Primers pRL423-VF and pRL423-VR were used for amplification from pRL415.

### NisB, NisC and NisPs purification

The protein NisPs was purified according to a previously described method (*22*). NisB and NisC were overexpressed and purified according to the previously described NisB purification method (*42*). Briefly, *E. coli* BL21 (DE3) cells were transformed with pET28-*nisB* and pET28-*nisC*. BL21 Rosetta (DE3) was transformed with pET28-*nisPs*. Several colony transformants were then grown in 50 mL of media supplemented with 50 μg/mL Kan at 37°C overnight. A 1% inoculation of a 2 L LB-antibiotic culture was grown aerobically at 37°C until OD_600_ reached 0.6–0.8. Then, 0.1 mM IPTG was added for NisPs induction and further grown for 3 h at 37°C. NisB and NisC cells were cooled to 18°C and IPTG was added to a final concentration of 0.5 mM (NisB) or 0.2 mM (NisC) and further grown for 20 h. For NisC overexpression, an additional 100 μM ZnCl_2_ was added to ensure enzymatic activity of NisC. After further growth, cells were harvested by centrifugation at 5,000 × *g* for 20 min at 4°C and resuspended in buffer A (20 mM Tris, pH 7.6, 500 mM NaCl, 10% glycerol). The cell suspension was lysed by homogenization at a variable pressure of 10,000–15,000 psig and centrifuged at 25,000 × *g* for 1 h at 4°C. The Ni-NTA column (GE Healthcare, Marlborough, MA, USA) was charged and washed with 2 column volumes (CV) of buffer A, and the filtered supernatant was applied to the column. The resin was washed with 2 CV each of buffers containing 0, 25, 50, 100, 200, and 500 mM imidazole. The purified protein was concentrated using Amicon Ultra-15 centrifugal filter devices (Millipore, Billerica, MA, USA) and the buffer was replaced with storage buffer (100 mM phosphate buffer, 10% glycerol, pH 7.6) via the PD-10 column (GE Healthcare). Protein concentrations were measured using a Pierce BCA Protein Assay Kit (Thermo Fisher Scientific) according to manufacturer’s instructions. Proteins were stored at −80°C after flash freezing in liquid nitrogen.

### CFPS reactions

CFPS reactions were performed to synthesize lanthipeptides. The previously described crude extract-based CFPS systems (*23*, *43*, *44*) was used for *in vitro* transcription and translation and supplemented with essential components for lanthipeptide PTMs. In the preparation of *E. coli* cell extracts, 1 mM IPTG was added during mid-log phase to induce T7 RNA polymerase overexpression, and cell lysate protein concentrations were measured using a Pierce BCA Protein Assay Kit (Thermo Fisher Scientific). The reaction mixture for CFPS consisted of the following components in a final volume of 8–400 μL containing 1.2 mM ATP; 0.85 mM each of GTP, UTP, and CTP; 34.0 μg/mL L-5-formyl-5,6,7,8-tetrahydrofolic acid (folinic acid); 170.0 μg/mL *E. coli* tRNA mixture; 130 mM potassium glutamate; 10 mM ammonium glutamate; 12 mM magnesium glutamate; 2 mM each of 20 natural amino acids; 0.33 mM nicotinamide adenine dinucleotide (NAD); 0.27 mM coenzyme-A (CoA); 1.5 mM spermidine; 1 mM putrescine; 4 mM sodium oxalate; 33 mM phosphoenolpyruvate (PEP); 10 μM ZnCl_2_, and 27% v/v cell extract. For each reaction, plasmids, purified NisB, and purified NisC were added at various concentrations. The CFPS reactions were performed at 30°C for 6 h. Reactions were terminated by incubation at 85°C for 10 min, and precipitated proteins were pelleted by centrifugation at 10,000 × *g* for 5 min. The resulting supernatant was subjected to downstream analyses.

### Qualitative and quantitative analyses by LC-MS-MS

Qualitative analyses of nisin Z and other lanthipeptides were performed using a previously described LC-MS-based method with modifications (*45*). Briefly, the supernatant of the CFPS reaction mixture or other solutions treated with trypsin were obtained after centrifugation at 10,000 × *g* for 10 min. Supernatants were desalted using Sep-Pak Vac C18 Cartridges (Waters, Milford, MA, USA) and subjected to LC-MS-MS. Chromatographic separation was performed using a Thermo Fisher Ultimate 3000 UPLC system equipped with a Thermo Fisher Hypersil GOLD C18 (2.1 × 100 mm, 3 μm) HPLC column; mobile phase A was H_2_O (0.1% formic acid) and mobile phase B was acetonitrile (ACN). The gradient program was (time, B%) 0 min, 10% B; 5 min, 10% B; 25 min, 95% B; 35 min, 95% B; 35.1 min, 10% B; 40 min, 10% B. The flow rate was 200 μL/min. The column temperature was 35°C and the injection volume was 10 μL. The sampler tray temperature was 8°C. Detection was performed using a Thermo Fisher Q Exactive Orbitrap MS with an ESI source in positive ion mode. Instrument parameters were as follows: sheath gas set to 35; auxiliary gas set to 5 (arbitrary units); spray voltage 3.5 kV; capillary temperature 320°C; probe heater temperature 250°C. Full scan trigger dd-MS_2_ mode was used for qualitative condition determination, and the settings were as follows: scan range 150–2,000 Da; MS resolution 70,000; MS_2_ resolution 17,500; isolation window 1.4 m/z; CE 30; dynamic exclusion of 2 s.

For lanthipeptides quantification, full scan mode was used. Nisin Z (eight-fold dehydrated component) was used to establish a standard curve. Angiotensin II was used as an internal standard, which was spiked at 10 ppb in standard curve working solutions. Additionally, 200 ppm BSA solution was used as a dilution reagent. The linear range was 10–20,000 ppb.

### Agar diffusion assay

The activity levels of nisin and new lanthipeptides were determined by a previously described agar diffusion method, with minor modifications (*46*, *47*). Briefly, a stock nisin solution (2000 IU/mL) was prepared by adding 50 mg of commercial nisin (10^6^ IU/g; Sigma-Aldrich, St. Louis, MO, USA) into 50 mL of sterile 0.02 mol/L HCl. Standard nisin solutions of 1000, 500, 250, 200, 100, 20, and 5 IU/mL were prepared using the stock solution and diluted with 0.02 mol/L HCl to construct a standard curve. The bioassay medium contained 1.2% tryptone, 0.75% yeast extract, 0.75% NaCl, 0.3% NaH_2_PO_4_, and 2% agar. After the addition of sterile 0.75% glucose and 0.5% Tween 20, the agar medium was cooled to 50°C and inoculated with 1.5% overnight culture of the indicator strain *M. luteus* NCIB 8166. The bioassay agar (25 mL) was aseptically poured into sterile Petri dishes, and several holes were made on each plate after solidification. Then, 2 μL of the supernatant of the CFPS mixture and an equal volume of the nisin Z standard solution were separately added to the holes. After incubation at 30°C for 18 h, a standard curve of the nisin zone of inhibition versus units of the nisin standard solution was created by measuring the zone diameter using digital calipers (TAJIMA Tool Co., Ltd., Shanghai, China) horizontally and vertically. Nisin concentrations for each CFPS mixture were estimated.

### In-silico prediction and selection of nisin analogs

Protein sequences for bacteria and fungi with lengths of 40 to 80 amino acid residues were retrieved from the NCBI database (accessed June 2018). The sequences containing the conserved motif of the core peptide (S×SLCTPGC×TG, where × denotes an arbitrary residue) were retained. The sequences were further reduced by the filter rule that LanBC-like or LanM-like proteins were found among the 10 genes in the upstream and downstream. Finally, after the exclusion of known peptides using BAGEL4 (*25*), 18 candidates were selected for experimental validation. These 18 sequences were adjusted to conform to the recognition site of NisP (if the N-terminal amino acid residues of the core peptide sequence were not “IT” or “VT,” “IT” was added to the N-terminus).

### Overexpression and purification of modified lanthipeptides precursor peptides

The overexpression and purification of His6-tagged mLanAs were performed as described previously (*5*). Briefly, overnight cultures were grown from a single recombinant *E. coli* colony transformant and used as inoculum to grow 1.5 L of Terrific Broth containing 50 mg/L Kan and 34 mg/L CmR at 37°C until the OD_600_ reached 0.6–0.8. The incubation temperature was adjusted to 18°C and the culture was induced with 0.5 mM IPTG. The induced cells were shaken continually at 18°C for an additional 18–20 h. The cells were harvested by centrifugation (11,900 × *g* for 10 min; Beckman JLA-10.500 rotor). The cell pellet was resuspended in 45 mL of start buffer (20 mM Tris, pH 8.0, 500 mM NaCl, 10% glycerol, containing a protease inhibitor cocktail from Roche Applied Science) and lysed by homogenization at variable pressures of 10,000–15,000 psig and centrifuged at 25,000 × *g* for 1 h at 4°C. The supernatant was loaded onto a nickel affinity column pre-equilibrated with start buffer. After loading, the column was washed with wash buffer (start buffer + 30 mM imidazole). The peptide was eluted from the column using elution buffer (start buffer + 500 mM imidazole), and then the elution buffer with the targeted modified peptide was replaced with PB buffer using an Amicon Ultra centrifugal filter (Millipore). Trypsin (2.5%) was added to the peptide solution and digested for 4 h at 37°C.

### N-Ethylmaleimide (NEM) alkylation assay of cyclization

NEM alkylation assays were performed according to a previously reported method (*7*). Briefly, an aliquot of the protease-digested peptide or full-length peptide solution was diluted into a two-fold volume of NEM alkylation buffer containing 500 mM HEPES, 3 mM NEM, 0.3 mM TCEP (pH 6.5). The reaction was incubated at 37°C for 30 min in the dark and analyzed by LC-MS-MS. Species containing uncyclized free Cys residues that were alkylated were identified by a mass increase of 125 Da for each adduct.

### Antibacterial assays

MICs were determined according to a previously described method (*48*), with minor modifications. Briefly, the test medium for clinical strain species was MHB broth (BD Difco), LB broth was used for *E. coli*, and bioassay medium containing 1.2% tryptone, 0.75% yeast extract, 0.75% NaCl, 0.3% NaH_2_PO_4_, and 0.75% glucose was used for *M. luteus*. Bacteria were grown overnight to the early stationary phase and adjusted in corresponding culture medium to 5.0 × 10^5^ CFU mL^−1^ in the wells of 96-well microtiter plates, mixed with varying concentrations of test compounds, and incubated at 37°C for 24 h. Cell growth was evaluated by measuring the optical density at 600 nm, and the MIC was defined as the lowest compound concentration at which no bacterial growth was observed.

### Target screening for nisin mutants with activity against E. coli

The reported nisin analogues [nisin A, nisin Z, nisin F (*27*), nisin Q (*28*), and nisin U (*29*)] were aligned and five amino acids at non-conserved sites (residues 4, 12, 15, 24, and 29 of nisin Z) were selected for saturation mutagenesis and inserted into pJL1 *Nde*I/*Bam*HI sites to form a mutant nisin precursor peptide library. The plasmid library was transformed into *E. coli*, and randomly selected monoclonal colonies were inoculated into 250 μL/well LB with 50 ug/mL Kan in 96-well plates and cultured at 37°C for 16 h; 50 μL of *E. coli* cultures were pipetted into an equal volume of 40% sterile glycerol for preservation. The remaining *E. coli* cultures were centrifuged at 4,500 rpm for 20 min. Then, 200 μL of sterile water was added to every well for cell resuspension and 0.5-mm glass beads were added at 0.24 g/well to suspended *E. coli* cells for lysis by shaking at 220 rpm for 90 min. The 96-well plate was centrifuged at 4,500 rpm for 5 min, 120 μL of the supernatant was pipetted into a new 96-well plate for cryogenic centrifugal concentration (1,500 rpm, −40°C, 1 pa), and 30 μL of sterile water was added to every well for residue resuspension. Then, the 96-well plate was centrifuged at 4,500 rpm for 1 min and 4 μL of the supernatant was pipetted into 5.5 μL of the CFPS system in a new 96-well plate, followed by incubation for 6 h at 30°C and 220 rpm. After incubation, the CFPS mixture in the 96-well plate was heated at 85°C for 5 min and centrifuged at 4,500 rpm for 10 min. Then, 2 μL of the supernatant was added to 5.0 × 10^5^ CFU mL^−1^ *E. coli* DH5α culture. OD_600_ was evaluated after co-cultivation at 37°C for 8 h.

After the first round of primary screening, the five mutant strains with the lowest OD_600_ values were selected from each 96-well plate as candidates. Another round of screening was performed using approximately 160 candidates among 3,000 mutants. Three wells were selected to set up the CFPS system using nisin Z as a control for rescreening. Therefore, after 8 h of the co-incubation of *E. coli* with the CFPS reaction system, the two mutants with the lowest ΔOD_600_ [ΔOD_600_ = OD_600_ (mutant) − OD_600_ (nisin Z)] were selected.

### Fed-batch fermentation of nisin Z production strains

The plasmids pRL415 and pRL423 were individually transformed into the J1-004 strain by electroporation as previously described (*49*) to generate the engineered strains RL405 and RL406, respectively. The industrial strain, J1-004, and colonies of engineered strains were incubated with seed medium overnight after cultivation on GM17 plates (w/v) (0.5% soy peptone, 0.5% beef extract, 0.5% tryptone, 0.25% yeast extract, 0.05% ascorbic acid, 0.025% MgSO_4_, 1.9% β-glycerophosphate disodium, 0.5% D-glucose g/L, 1.8% agar) at 30°C for 48 h. The seed medium (w/v) contained 1.5% peptone, 1.5% yeast extract, 1.5% sucrose, 2.0% KH_2_PO_4_, 0.15% NaCl, 0.3% corn steep liquor, 0.26% cysteine, and 0.015% MgSO_4_7H_2_O. Then 5% of the seed culture was inoculated into 3 L of fermentation medium in a 7-L fermenter and 10 mol/L NaOH was used to adjust the medium pH to 7.2. The fermentation medium contained yeast extract (1%), sucrose (0.6%), KH_2_PO_4_ (0.5%), NaCl (0.1%), corn steep liquor (3%), cysteine (0.26%), and MgSO_4_·7H_2_O (0.015%). The pH of the fermentation broth was maintained at 6.7 by the addition of 10 mol/L NaOH. A sucrose solution (50%, w/v) was used to maintain the residual sugar concentration in the fermentation broth at approximately 1% from 3 h-13 h. Samples broth (10 mL) were removed after 16 h of fermentation. After centrifugation, the supernatant was subjected to a quantitative bioassay to determine nisin production.

### Statistical analysis

For quantification of lanthipeptides’ titer in CFPS experiments, flask fermentation, and fed-batch fermentation, experiments were repeated three times independently and quantified by agar diffusion assay. Data shown are mean ±SD from the three replicates. In agar diffusion assays, every sample was repeated three times independently for measurement, the average value was calculated as the result of these independent experiments. For MIC assays of *M. luteus*, *E. faecalis*, *S. aureus*, and MRSA, experiments were repeated three times independently. For OD_600_ detection of co-culture lanthipeptides with *E. coli*, experiments were repeated three times independently. For screening nisin mutant library in CFPS platform, experiments were performed once unless otherwise indicated.

## Supporting information

manuscript

## Author contributions

R.L., Yuchen Z. and T.L. designed the study. R.L., Yuchen Z., G.Z., S.F., and Y.X. performed all experiments, B.H. provided bioinformatics analysis of potential lanthipeptides. All authors analyzed the data. R.L. and T.L. wrote the manuscript.

## Data and materials availability

All data needed to evaluate the conclusions in the paper are present in the paper and/or the Supplementary Materials. Additional data related to this paper may be requested from the authors.

## References

1. P. G. Arnison, M. J. Bibb, G. Bierbaum, A. A. Bowers, T. S. Bugni, G. Bulaj, J. A. Camarero, D. J. Campopiano, G. L. Challis, J. Clardy, P. D. Cotter, D. J. Craik, M. Dawson, E. Dittmann, S. Donadio, P. C. Dorrestein, K. D. Entian, M. A. Fischbach, J. S. Garavelli, U. Goransson, C. W. Gruber, D. H. Haft, T. K. Hemscheidt, C. Hertweck, C. Hill, A. R. Horswill, M. Jaspars, W. L. Kelly, J. P. Klinman, O. P. Kuipers, A. J. Link, W. Liu, M. A. Marahiel, D. A. Mitchell, G. N. Moll, B. S. Moore, R. Muller, S. K. Nair, I. F. Nes, G. E. Norris, B. M. Olivera, H. Onaka, M. L. Patchett, J. Piel, M. J. Reaney, S. Rebuffat, R. P. Ross, H. G. Sahl, E. W. Schmidt, M. E. Selsted, K. Severinov, B. Shen, K. Sivonen, L. Smith, T. Stein, R. D. Sussmuth, J. R. Tagg, G. L. Tang, A. W. Truman, J. C. Vederas, C. T. Walsh, J. D. Walton, S. C. Wenzel, J. M. Willey, W. A. van der Donk, Ribosomally synthesized and post-translationally modified peptide natural products: overview and recommendations for a universal nomenclature. Nat Prod Rep 30, 108–160 (2013).

2. L. E. Cooper, B. Li, W. A. van der Donk, Biosynthesis and mode of action of lantibiotics. Chem Rev 105, 633–684 (2005).

3. M. A. Skinnider, C. W. Johnston, R. E. Edgar, C. A. Dejong, N. J. Merwin, P. N. Rees, N. A. Magarvey, Genomic charting of ribosomally synthesized natural product chemical space facilitates targeted mining. Proc Natl Acad Sci U S A 113, E6343–E6351 (2016).

4. J. I. Tietz, C. J. Schwalen, P. S. Patel, T. Maxson, P. M. Blair, H. C. Tai, U. I. Zakai, D. A. Mitchell, A new genome-mining tool redefines the lasso peptide biosynthetic landscape. Nat Chem Biol 13, 470–478 (2017).

5. Y. Shi, X. Yang, N. Garg, W. A. van der Donk, Production of lantipeptides in Escherichia coli. J Am Chem Soc 133, 2338–2341 (2011).

6. A. J. van Heel, T. G. Kloosterman, M. Montalban-Lopez, J. Deng, A. Plat, B. Baudu, D. Hendriks, G. N. Moll, O. P. Kuipers, Discovery, production and modification of five novel lantibiotics using the promiscuous nisin modification machinery. ACS Synth Biol 5, 1146–1154 (2016).

7. X. Yang, K. R. Lennard, C. He, M. C. Walker, A. T. Ball, C. Doigneaux, A. Tavassoli, W. A. van der Donk, A lanthipeptide library used to identify a protein-protein interaction inhibitor. Nat Chem Biol 14, 375–380 (2018).

8. S. Schmitt, M. Montalban-Lopez, D. Peterhoff, J. Deng, R. Wagner, M. Held, O. P. Kuipers, S. Panke, Analysis of modular bioengineered antimicrobial lanthipeptides at nanoliter scale. Nat Chem Biol 15, 437–443 (2019).

9. T. Si, Q. Tian, Y. Min, L. Zhang, J. V. Sweedler, W. A. van der Donk, H. Zhao, Rapid screening of lanthipeptide analogs via in-colony removal of leader peptides in Escherichia coli. J Am Chem Soc 140, 11884–11888 (2018).

10. G. A. Hudson, Z. Zhang, J. I. Tietz, D. A. Mitchell, W. A. van der Donk, In vitro biosynthesis of the core scaffold of the thiopeptide thiomuracin. J Am Chem Soc 137, 16012–16015 (2015).

11. J. M. Shin, J. W. Gwak, P. Kamarajan, J. C. Fenno, A. H. Rickard, Y. L. Kapila, Biomedical applications of nisin. J Appl Microbiol 120, 1449–1465 (2016).

12. P. D. Cotter, C. Hill, R. P. Ross, Bacteriocins: developing innate immunity for food. Nat Rev Microbiol 3, 777–788 (2005).

13. L. Devuyst, Nisin production variability between natural Lactococcus lactis subsp lactis strains. Biotechnol Lett 16, 287–292 (1994).

14. T. V. Polyudova, L. M. Lemkina, G. N. Likhatskaya, V. P. Korobov, Optimization of production conditions and 3D-structure modeling of novel antibacterial peptide of lantibiotic family. Appl Biochem Microbiol 53, 40–46 (2017).

15. P. D. Cotter, L. A. Draper, E. M. Lawton, O. McAuliffe, C. Hill, R. P. Ross, Overproduction of wild-type and bioengineered derivatives of the lantibiotic lacticin 3147. Appl Environ Microbiol 72, 4492–4496 (2006).

16. C. E. Hodgman, M. C. Jewett, Cell-free synthetic biology: thinking outside the cell. Metab Eng 14, 261–269 (2012).

17. E. D. Carlson, R. Gan, C. E. Hodgman, M. C. Jewett, Cell-free protein synthesis: applications come of age. Biotechnol Adv 30, 1185–1194 (2012).

18. F. Cheng, T. M. Takala, P. E. Saris, Nisin biosynthesis in vitro. J Mol Microbiol Biotechnol 13, 248–254 (2007).

19. S. R. Fleming, T. E. Bartges, A. A. Vinogradov, C. L. Kirkpatrick, Y. Goto, H. Suga, L. M. Hicks, A. A. Bowers, Flexizyme-enabled benchtop biosynthesis of thiopeptides. J Am Chem Soc 141, 758–762 (2019).

20. T. Ozaki, K. Yamashita, Y. Goto, M. Shimomura, S. Hayashi, S. Asamizu, Y. Sugai, H. Ikeda, H. Suga, H. Onaka, Dissection of goadsporin biosynthesis by in vitro reconstitution leading to designer analogues expressed in vivo. Nat Commun 8, 14207 (2017).

21. R. Khusainov, G. N. Moll, O. P. Kuipers, Identification of distinct nisin leader peptide regions that determine interactions with the modification enzymes NisB and NisC. FEBS Open Bio 3, 237–242 (2013).

22. M. Lagedroste, S. H. J. Smits, L. Schmitt, Substrate specificity of the secreted nisin leader peptidase NisP. Biochemistry 56, 4005–4014 (2017).

23. Y. Zhang, H. Qianyin, D. Zixin, X. Yancheng, L. Tiangang, Enhancing the efficiency of cell-free protein synthesis system by systematic titration of transcription and translation components. Biochem Eng J, 138, 47–53 (2018).

24. T. C. Penna, A. F. Jozala, L. C. De Lencastre Novaes, A. Pessoa, O. Cholewa, Production of nisin by Lactococcus lactis in media with skimmed milk. Appl Biochem Biotechnol 121-124, 619–637 (2005).

25. A. J. van Heel, A. de Jong, C. Song, J. H. Viel, J. Kok, O. P. Kuipers, BAGEL4: a user-friendly web server to thoroughly mine RiPPs and bacteriocins. Nucleic Acids Res 46, W278–W281 (2018).

26. D. Field, M. Begley, P. M. O’Connor, K. M. Daly, F. Hugenholtz, P. D. Cotter, C. Hill, R. P. Ross, Bioengineered nisin A derivatives with enhanced activity against both Gram positive and Gram negative pathogens. PLoS One 7, e46884 (2012).

27. M. de Kwaadsteniet, K. Ten Doeschate, L. M. Dicks, Characterization of the structural gene encoding nisin F, a new lantibiotic produced by a Lactococcus lactis subsp. lactis isolate from freshwater catfish (Clarias gariepinus). Appl Environ Microbiol 74, 547–549 (2008).

28. T. Zendo, M. Fukao, K. Ueda, T. Higuchi, J. Nakayama, K. Sonomoto, Identification of the lantibiotic nisin Q, a new natural nisin variant produced by Lactococcus lactis 61-14 isolated from a river in Japan. Biosci Biotechnol Biochem 67, 1616–1619 (2003).

29. R. E. Wirawan, N. A. Klesse, R. W. Jack, J. R. Tagg, Molecular and genetic characterization of a novel nisin variant produced by Streptococcus uberis. Appl Environ Microbiol 72, 1148–1156 (2006).

30. L. Zhou, A. J. van Heel, M. Montalban-Lopez, O. P. Kuipers, Potentiating the activity of nisin against Escherichia coli. Front Cell Dev Biol 4, 7 (2016).

31. Q. Li, M. Montalban-Lopez, O. P. Kuipers, Increasing the antimicrobial activity of nisin-based lantibiotics against Gram-negative pathogens. Appl Environ Microbiol 84, (2018).

32. J. Garamella, R. Marshall, M. Rustad, V. Noireaux, The all E. coli TX-TL toolbox 2.0: a platform for cell-free synthetic biology. ACS Synth Biol 5, 344–355 (2016).

33. F. Caschera, V. Noireaux, Synthesis of 2.3 mg/ml of protein with an all Escherichia coli cell-free transcription-translation system. Biochimie 99, 162–168 (2014).

34. K. A. Calhoun, J. R. Swartz, An economical method for cell-free protein synthesis using glucose and nucleoside monophosphates. Biotechnol Prog 21, 1146–1153 (2005).

35. Q. Cai, J. A. Hanson, A. R. Steiner, C. Tran, M. R. Masikat, R. Chen, J. F. Zawada, A. K. Sato, T. J. Hallam, G. Yin, A simplified and robust protocol for immunoglobulin expression in Escherichia coli cell-free protein synthesis systems. Biotechnol Prog 31, 823–831 (2015).

36. A. J. van Heel, D. Mu, M. Montalban-Lopez, D. Hendriks, O. P. Kuipers, Designing and producing modified, new-to-nature peptides with antimicrobial activity by use of a combination of various lantibiotic modification enzymes. ACS Synth Biol 2, 397–404 (2013).

37. B. J. Burkhart, N. Kakkar, G. A. Hudson, W. A. van der Donk, D. A. Mitchell, Chimeric leader peptides for the generation of non-natural hybrid RiPP products. ACS Cent Sci 3, 629–638 (2017).

38. S. J. Moore, J. T. MacDonald, S. Wienecke, A. Ishwarbhai, A. Tsipa, R. Aw, N. Kylilis, D. J. Bell, D. W. McClymont, K. Jensen, K. M. Polizzi, R. Biedendieck, P. S. Freemont, Rapid acquisition and model-based analysis of cell-free transcription-translation reactions from nonmodel bacteria. Proc Natl Acad Sci U S A 115, E4340–E4349 (2018).

39. A. S. Karim, M. C. Jewett, A cell-free framework for rapid biosynthetic pathway prototyping and enzyme discovery. Metab Eng 36, 116–126 (2016).

40. D. G. Gibson, Enzymatic assembly of overlapping DNA fragments. Methods Enzymol 498, 349–361 (2011).

41. J. Lubelski, R. Rink, R. Khusainov, G. Moll, O. Kuipers, Biosynthesis, immunity, regulation, mode of action and engineering of the model lantibiotic nisin. Cellular and Molecular Life Sciences 65, 455–476 (2008).

42. M. A. Ortega, Y. Hao, Q. Zhang, M. C. Walker, W. A. van der Donk, S. K. Nair, Structure and mechanism of the tRNA-dependent lantibiotic dehydratase NisB. Nature 517, 509–+ (2015).

43. Y. C. Kwon, M. C. Jewett, High-throughput preparation methods of crude extract for robust cell-free protein synthesis. Sci Rep 5, 8663 (2015).

44. M. C. Jewett, J. R. Swartz, Mimicking the Escherichia coli cytoplasmic environment activates long-lived and efficient cell-free protein synthesis. Biotechnol Bioeng 86, 19–26 (2004).

45. N. Schneider, K. Werkmeister, M. Pischetsrieder, Analysis of nisin A, nisin Z and their degradation products by LCMS/MS. Food Chem 127, 847–854 (2011).

46. T. Pongtharangkul, A. Demirci, Evaluation of agar diffusion bioassay for nisin quantification. Appl microbiol biotechnol 65, 268–272 (2004).

47. T. J. Oman, W. A. Van Der Donk, Insights into the mode of action of the two-peptide lantibiotic haloduracin. ACS chem bio 4, 865–874 (2009).

48. Y. X. Li, Z. Zhong, W. P. Zhang, P. Y. Qian, Discovery of cationic nonribosomal peptides as Gram-negative antibiotics through global genome mining. Nat Commun 9, (2018).

49. H. Holo, I. F. Nes, High-frequency transformation, by electroporation, of Lactococcus lactis subsp. cremoris grown with glycine in osmotically stabilized media. Appl Environ Microbiol 55, 3119–3123 (1989).

