## Supplementary material for "A cell-free platform based on nisin biosynthesis for discovering novel lanthipeptides and guiding their overproduction *in vivo*": manuscript

### Supplementary Information

#### 1. Supplementary Materials and Methods

*Semi-preparative HPLC separation of commercialized nisin A*

Purification was performed using the UltiMate3000 Semi-preparative HPLC (Thermo Fisher). The system was equipped with a UV detector and UltiMate 3000 Fraction Collector. A Fisher Hypersil Gold column (250 × 10 mm, i.d., 5 μm) was employed for peptide separation at 35 °C. The injection volume was 100 μL. The sampler tray temperature was 10 °C. The column flow rate was 2 mL/min, and detection was set at 215 nm. The column was used under the following conditions: mobile phase A was $H_2O$ (0.1% formic acid); mobile phase B was acetonitrile (ACN). The gradient program was (time, B%) 0 min, 5% B; 1 min, 5% B; 30 min, 90% B; 35 min, 90% B; 36 min, 5% B; 50 min, 5% B. All chromatographic peaks were collected during the preliminary experiment, and all fractions were detected by LC-MS-MS. The desired factions were then collected.

*LC-MS analysis of nisin Z and targeted proteomics analysis*

The original strain, J1-004, and the engineered *L. lactis* cells collected from the fermentation broth were analyzed by LC-MS for nisin Z and by a targeted proteomics approach according to a previously reported method (*1*). Briefly, *L. lactis* cells collected from the fermentation broth were pelleted by centrifugation 8,000 × *g* for 10 min at 4 °C, collected, and washed three times with wash buffer (100 mM NaCl, 25 mM Tris−HCl, pH 7.5). The wet cell pellet was suspended in an equal volume (1 g wet cell weight/1 mL of buffer) of lysis buffer (8 M urea, 2 M thiourea, 75 mM NaCl, 4% (w/v) 3-[(3-cholamidopropyl)-dimethylammonio]-1-propanesulfonate (CHAPS), 50 mM Tris−HCl (pH 8.0) and 1 complete EDTA-Free protease inhibitor cocktail tablet (Roche, Indianapolis, IN, USA) per 10 mL of buffer). The suspended sample was vortexed for 30 s twice and disrupted by sonication (*2*). The supernatant from the lysed cells was collected by centrifugation (13,000 × *g* for 45 min at 4 °C). Proteins from the cell lysates were measured using a noninterference protein assay kit (Sangon Biotech, Shanghai, China) and adjusted to 2 μg/μL using lysis buffer. First, 50 μL of the supernatant (100 μg of total protein) was mixed with an equal volume of 100 mM ammonium bicarbonate buffer (pH 8.0). Next, the sample was reduced at 30 °C for 1 h by the addition of 3 mM tris (2-carboxyethyl)-phosphine (TCEP) and alkylated by the addition of 15 mM iodoacetamide (IAA). The samples were incubated in dark conditions at 30 °C for an additional 1 h. The sample was diluted with ammonium bicarbonate buffer to reduce the urea concentration to 1 M. Trypsin was added to the mixture (trypsin/total protein 1:50, w/w) and incubated at 37 °C for 14 h. The detergent and salt in the digested peptide sample were removed by passage through a Pierce Detergent Removal Spin Column (Thermo Fisher Scientific) and a SepPak C18 cartridge (Waters Corp.), respectively. The purified peptides were freeze-dried and stored at -80 °C for subsequent liquid chromatography-tandem mass spectrometry (LC-MS-MS) analysis.

The peptide samples were analyzed using a hybrid quadrupole-time-of-flight (TOF) liquid chromatography (LC) tandem mass (MS/MS) spectrometer (TripleTOF 5600+, AB Sciex, Foster City, CA, USA) equipped with a nanospray ion source. Peptides were first loaded onto a C18 trap column (5 μm, 5 × 0.3 mm; Agilent Technologies, Santa Clara, CA, USA) and then eluted into a C18 analytical column (75 μm × 150 mm, 3 μm particle size, 100 Å pore size; Eksigent, Dublin, CA, USA). Mobile phase A (3% DMSO, 97% $H_2O$, 0.1% formic acid) and mobile phase B (3% DMSO, 97% ACN, 0.1% formic acid) were used to establish a 100-min gradient as follows: 0 min of 5% B, 65 min of 5–23% B, 20 min of 23–52% B, 1 min of 52–80% B, maintenance at 80% B for 4 min, 0.1 min of 80–5% B, and a final step of 5% B for 10 min. A constant flow rate was set at 300 nL/min. MS scans were conducted from 350 amu-1500 amu, with a 250-ms time span. For the MS/MS analysis, each scan cycle consisted of one full-scan mass spectrum (with m/z ranging from 350-1500 and charge states from 2-5) followed by 40 MS/MS events. The threshold count was set to 120 to activate MS/MS accumulation and former target ion exclusion was set to 18 s. Raw data obtained by TripleTOF 5600+ were analyzed using ProteinPilot 5.0 (AB SCIEX) against the designated proteome database.

#### 2. Supplementary Figures

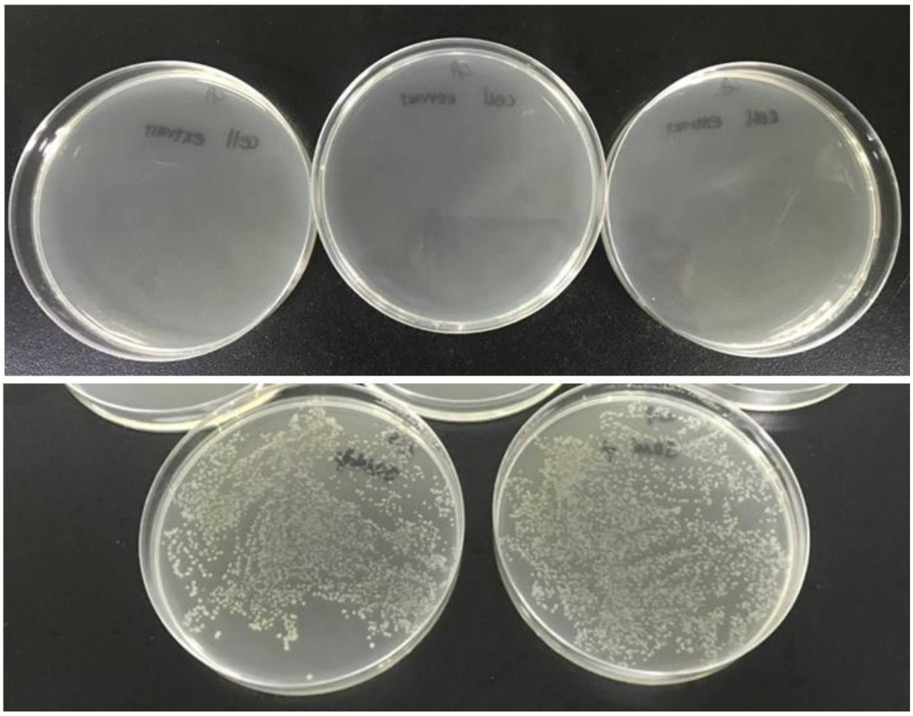

**Fig. S1 | Cytoplasmic extract control plates**

All LB agar (no antibiotic) plates were incubated at 37 °C for 16 h and 27 % v/v of the cell extract was used for each CFPS reaction. **Top:** Approximately 50 µL of *E. coli* cell extracts was plated on each of the three plates. No *E. coli* colonies grew on either plate. **Bottom:** Approximately 50 µL of *E. coli* ($5.0 \times 10^5$ CFU $mL^{-1}$) cells were added to the plate as a positive control.

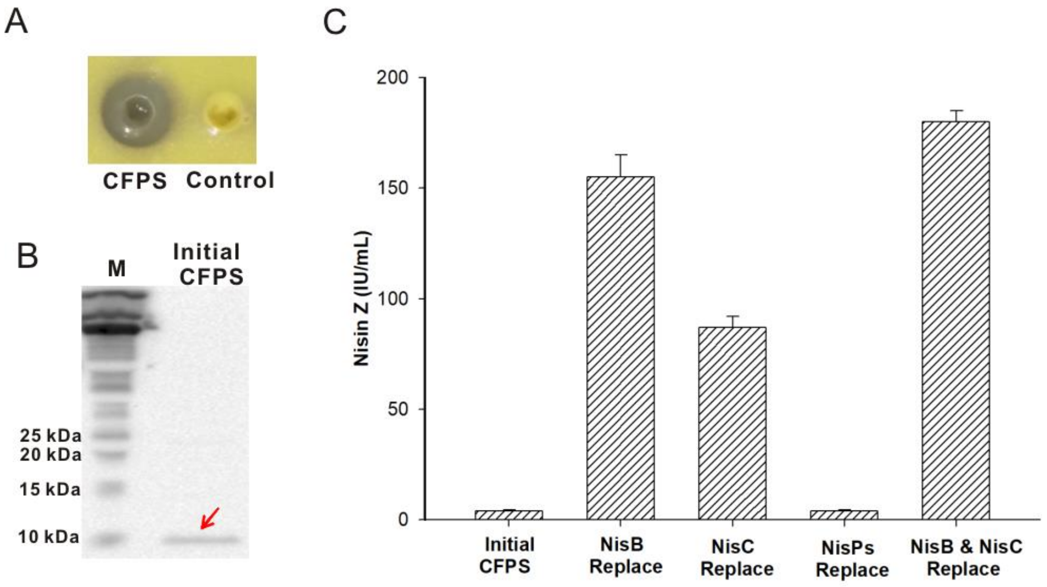


**Fig. S2 | Analysis of the nisin CFPS system**

**(A)** Validation of antimicrobial activity by the agar diffusion assay. An obvious zone of inhibition was detected on the *M. luteus* plate using the nisin CFPS reaction mixture (CFPS). Control: CFPS reaction without pJL1-*nisZ*, pET28a-*nisB*, pET28a-*nisC*, and pET28a-*nisP*. (**B**) Western blot analysis of expressed proteins (NisZ, NisB, NisC, and NisP) involved in Nisin CFPS. Red arrow indicates the partially modified Nisin precursor with an N-terminal His6-tag. A primary anti-His6 tag mouse monoclonal antibody was diluted 1:2,000 in blocking buffer (2% nonfat milk in TBST) before use. Horseradish peroxidase (HRP)-conjugated goat anti-mouse IgG was used as the secondary antibody, and the SuperSignal West Pico Plus Chemiluminescent Substrate was used to visualize proteins via chemiluminescence. (**C**) Replacement of poorly expressed protein encoding plasmids with purified enzymes in CFPS. The final concentrations of purified proteins were fixed at 500 nM.

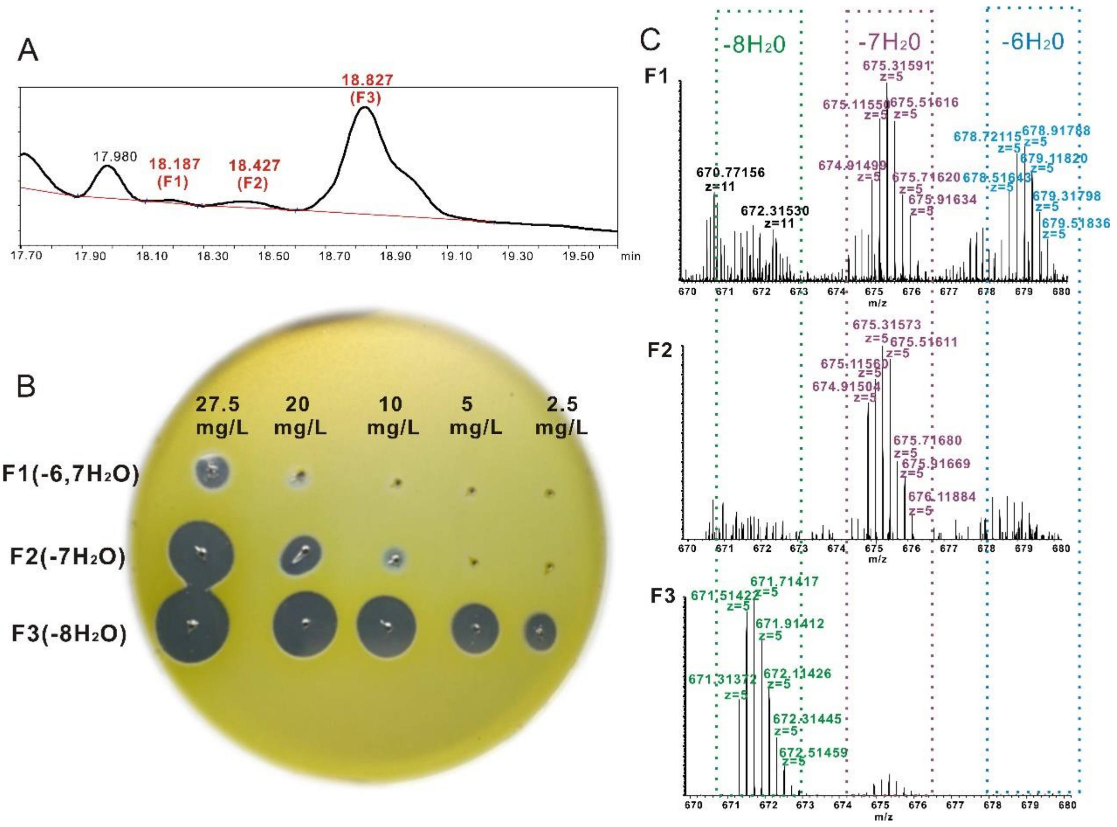


**Fig. S3 | Identification of the most efficient nisin A component**

(**A**) Separation of different components of nisin A standard by semi-preparative HPLC. F1, F2, and F3 were collected. (**B**) Antibacterial activity assay of different components using the agar diffusion method. The concentrations of F1, F2, and F3 were set to 2.5 mg/L, 5 mg/L, 10 mg/L, 20 mg/L, and 27.5 mg/L, and 2 μL of each sample was used for the antibacterial activity test. (**C**) High-resolution mass spectrometry of different components. The F1 component consists of dehydrated (-6, -7$H_2O$) nisin A, the F2 component consists of dehydrated (-7 $H_2O$) nisin A, and the F3 component consists of dehydrated (-8 $H_2O$) nisin A.

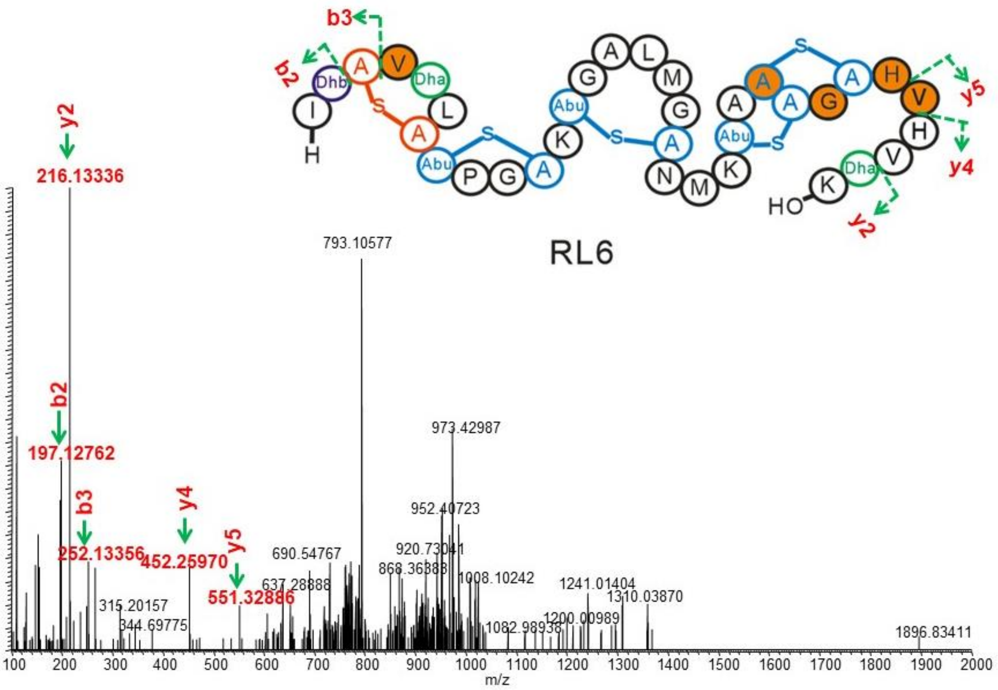


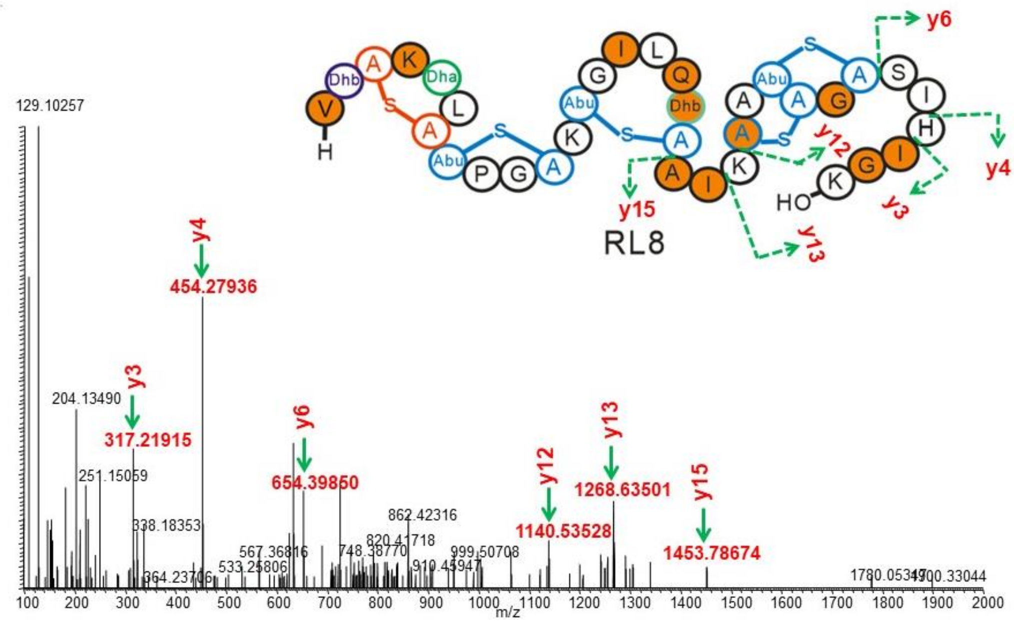


**Fig. S4.** Continued on next page.

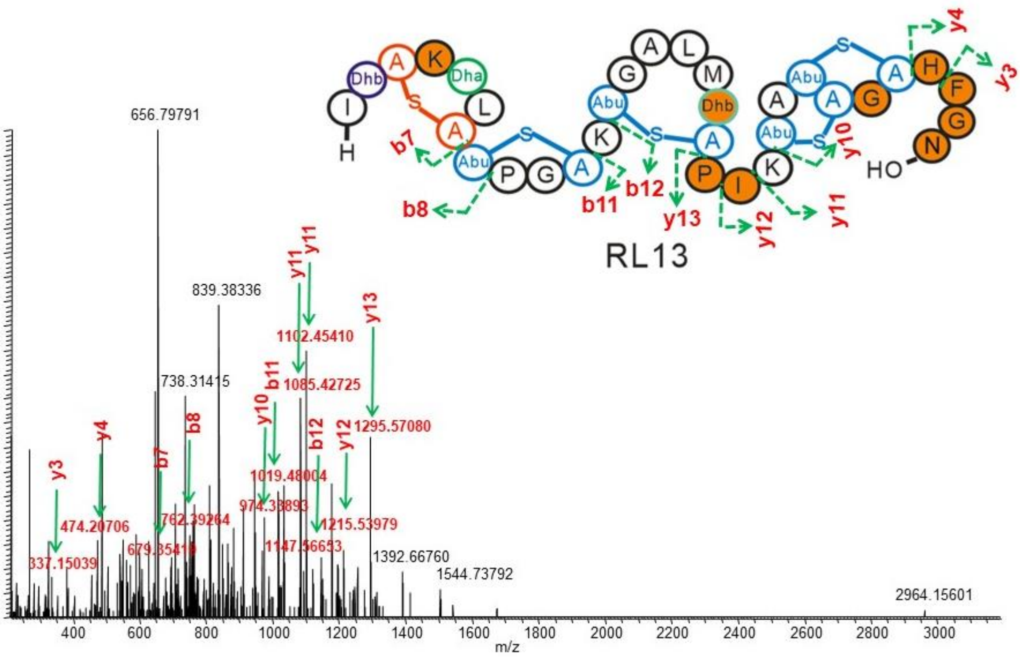


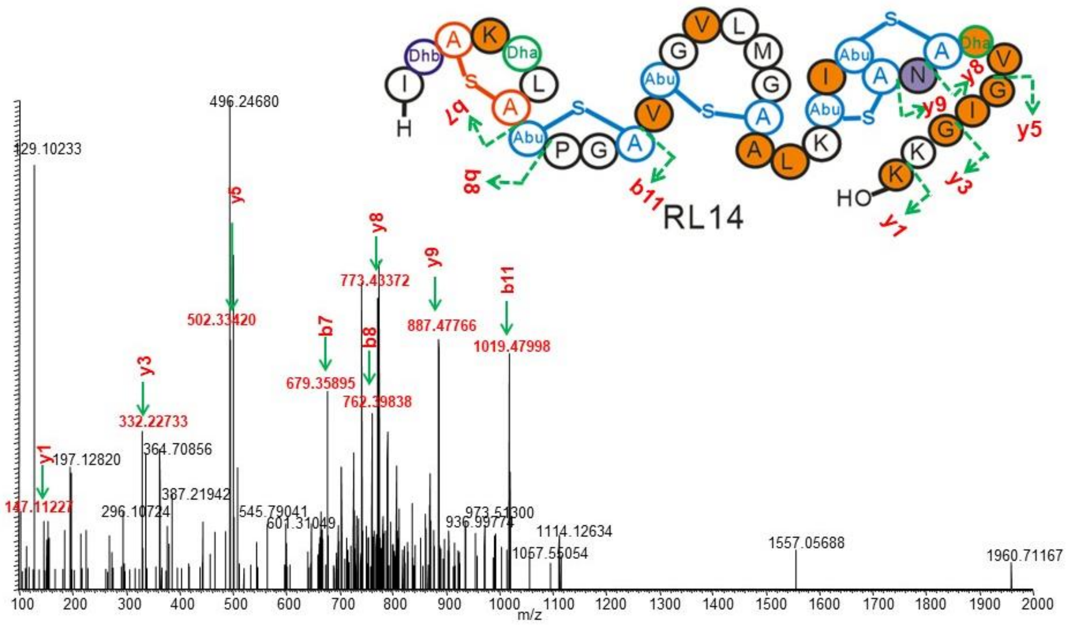


**Fig. S4.** Continued on next page.

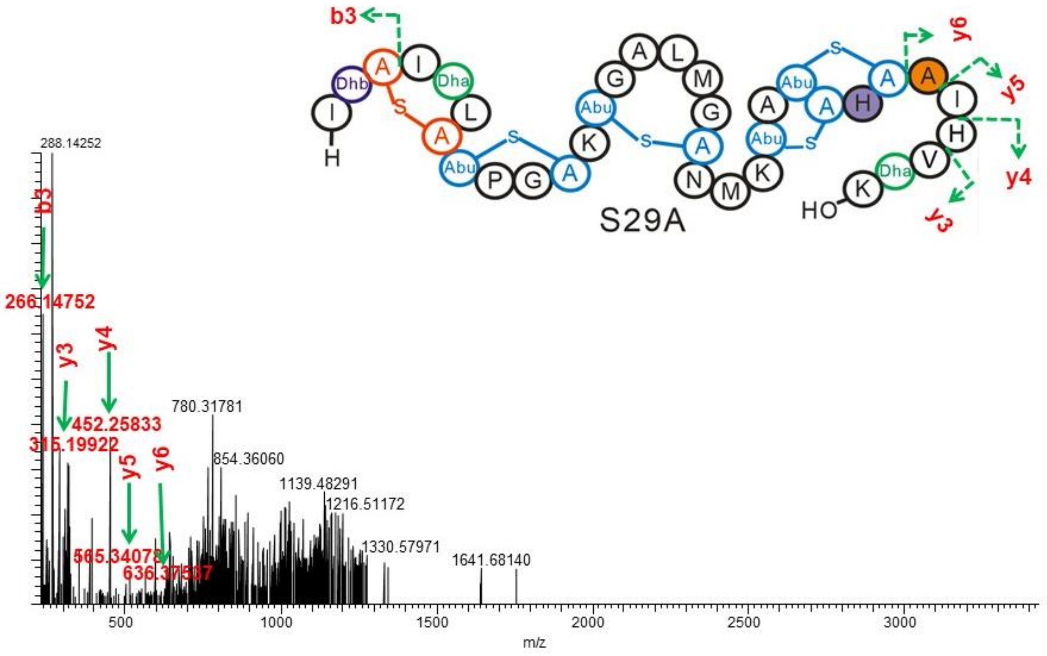


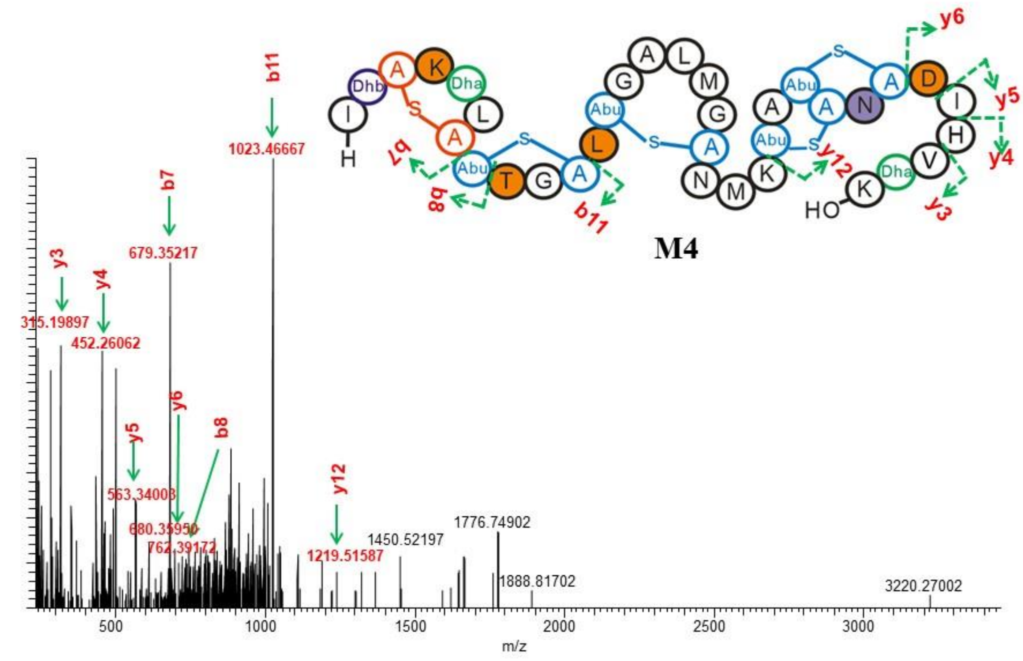


**Fig. S4.** Continued on next page.

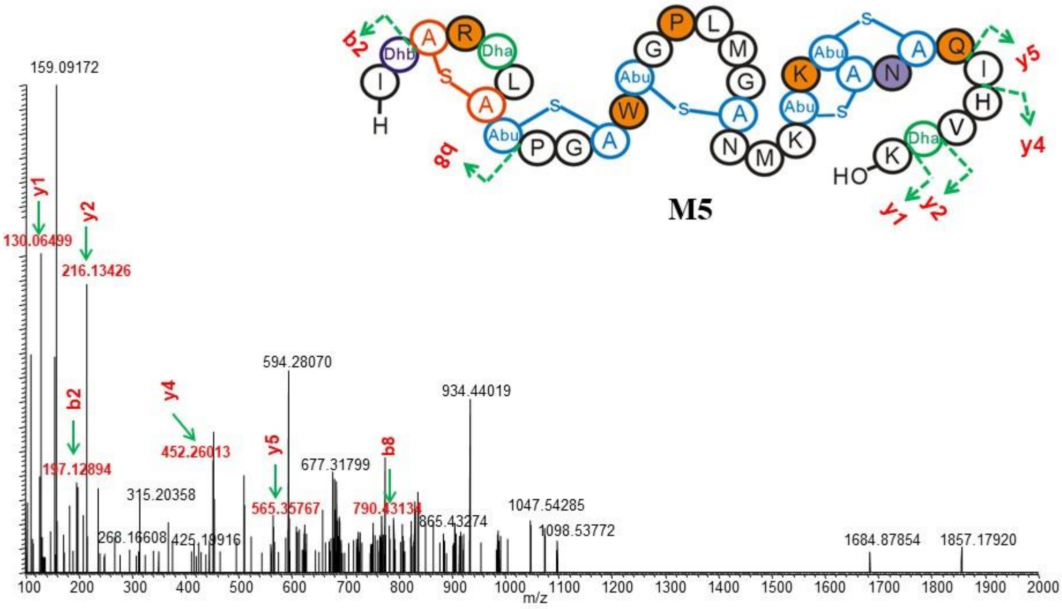


**Fig. S4 | Structure of nisin analogs and nisin mutants determined by high-resolution mass spectrometry**

Brown circle markers represent the amino acid residues that are different from nisin residues. Dha: dehydrolalanine; Dhb: dehydrobutyrine; Abu: 2-aminobutyric acid.

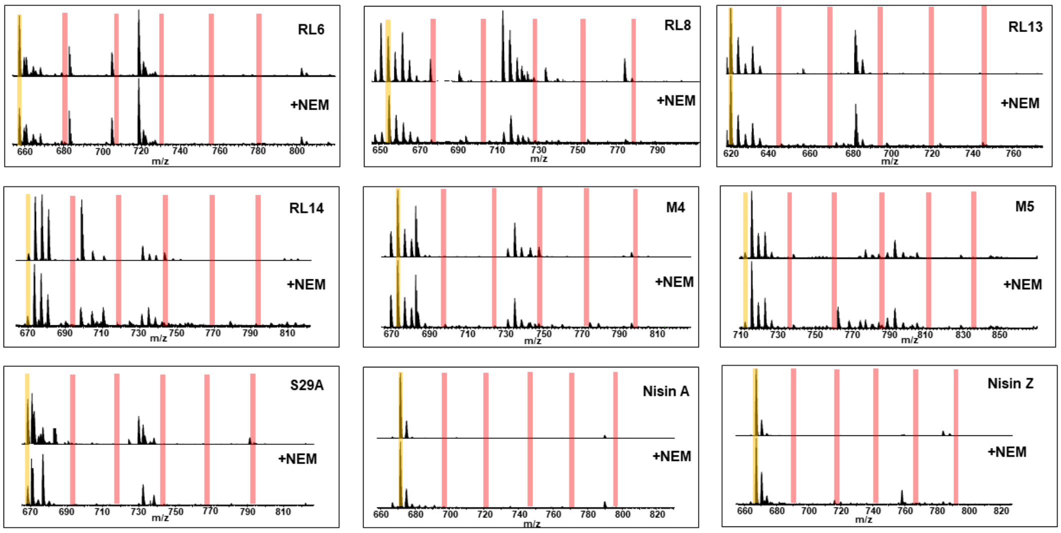


**Fig. S5 | Summary of the LC-MS analysis of lanthipeptides studied in this research**

All ions are in the +5-charge state. Purified modified precursor peptides were digested with trypsin to form mature RL6, RL8, RL13, RL14, M4, M5, and S29A and treated with NEM. Commercialized nisin A and nisin Z were also treated with NEM. The eight-fold dehydrated core peptides are highlighted in yellow.

Alkylated core peptides with 1–5 NEM adducts are highlighted in red. All five thioester rings were formed in mature lanthipeptides.

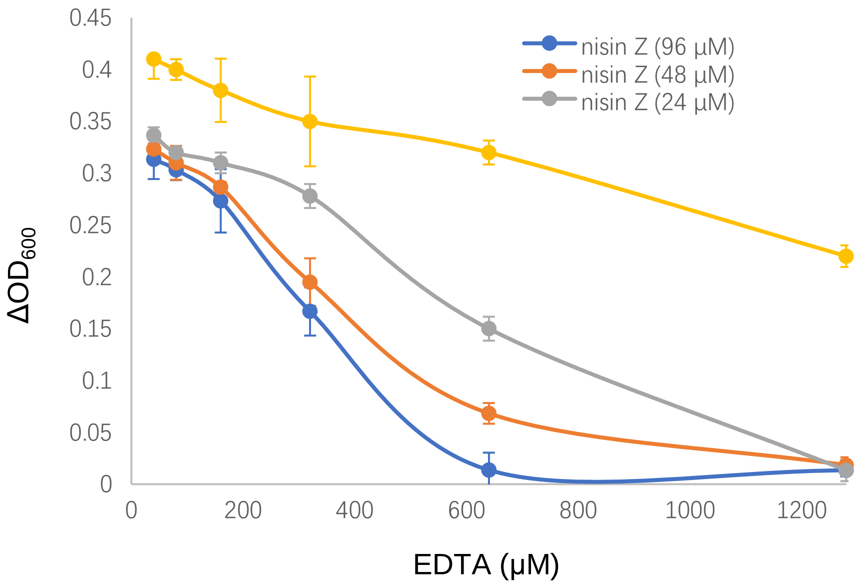


**Fig. S6 | $OD_{600}$ of *E. coli* DH5α in LB medium with different concentrations of EDTA and Nisin after 18 h of co-culture**

$\Delta OD_{600}$ indicates the difference between readings with different concentrations of blank media. Previous studies have reported that nisin has a significant inhibitory effect on *E. coli (3-5)*, however, uncommon *E. coli* strains were used. In the reported 20-fold nisin concentration condition, DH5α did not exhibit complete inhibition. To enable comparisons with previous studies, EDTA was added to the *E. coli* culture to increase the sensitivity of *E. coli* to nisin. After 18 h, *E. coli* DH5α growth was only slightly inhibited following addition of 320 μM EDTA, however, the inhibitory effects of various concentrations of nisin on *E. coli* were obvious. Error bars are based on three independent replicates.

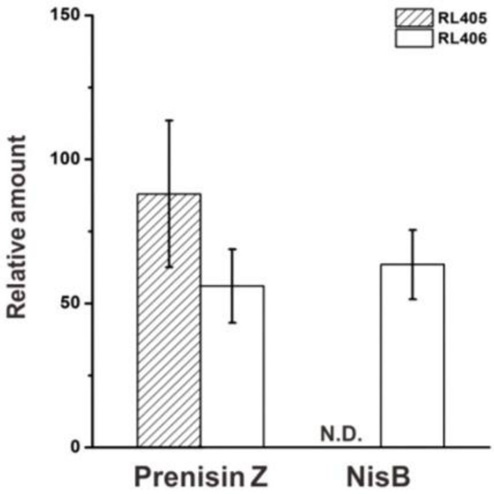


**Fig. S7 | Proteomic analysis of targeted proteins involved in nisin biosynthesis in engineered strains**

Relative amount: number of identified specific peptides with 95% confidence; N.D.: Not detected. Error bars are based on three independent replicates.

#### 2. Supplementary Tables

**Table S1. Strains used in this research**

| Strains | Describe | Source |
|---|---|---|
| DH10β | Plasmid construction | Our Lab Preserve |
| XL1-Blue | Plasmid construction | Our Lab Preserve |
| BL21(DE3) | Protein/ mLanAs overexpression. | Our Lab Preserve |
| BL21Rosseta(DE3) | NisPs overexpression. | Our Lab Preserve |
| *Micrococcus luteus* | Indicator strain for nisin antibacterial activity test | NCIB 8166 |
| BL21(DE3)/pYZ95/pYX126 | Expression of sumo-tagged mRL6 | This study |
| BL21(DE3)/pYZ96/pYX126 | Expression of sumo-tagged mRL8 | This study |
| BL21(DE3)/pYX122/pYX125 | Expression of sumo-tagged mRL13 | This study |
| BL21(DE3)/pYX123/pYX125 | Expression of sumo-tagged mRL14 | This study |
| BL21(DE3)/pYZ92/pYX126 | Expression of sumo-tagged mM4 | This study |
| BL21(DE3)/pYZ93/pYX126 | Expression of sumo-tagged mM5 | This study |
| BL21(DE3)/pYZ97/pYX126 | Expression of sumo-tagged S29A | This study |
| *Enterococcus faecalis* | Clinical standard strain for antibacterial testing | ATCC 29212 |
| *Staphylococcus aureus* | Clinical standard strain for antibacterial testing | ATCC 25923 |
| Methicillin-resistant *Staphylococcus aureus* (MRSA) | Clinical isolation of antibiotic test resistant strains | Renmin hospital of Wuhan University |
| *Lactococcus lactis* J1-004 | Nisin Z industrial producing strain | J1 Biotech. Co. |
| RL405 | J1-004/pRL415, Overexpress *nisZ* in J1-004 | This study |
| RL406 | J1-004/pRL423, Overexpress nisZ and *nisB* in J1-004 | This study |
| BL21(DE3)/ pYX106 | Expression of His6-tagged mRL6 (without *nisB* overexpressed) in *E. coli* | This study |
| BL21(DE3)/ pYX106//pYX125 | overexpress His6-tagged mRL6 (with *nisB* overexpressed) in *E. coli* | This study |

**Table S2 MIC values of several lanthipeptides against microorganisms**

| MIC* mg/L(μM) | *M. luteus* | *E. faecalis* | *S. aureus* | MRSA |
|---|---|---|---|---|
| Nisin A | 0.06 (0.018) | 250 (74.53) | 125 (37.27) | 125 (37.27) |
| Nisin Z | 0.12 (0.036) | 250 (75.05) | 250 (75.05) | 125 (37.52) |
| RL6 | 0.12 (0.037) | 250 (76.17) | >250 (>76.17) | >250 (>76.17) |
| RL8 | 0.24 (0.073) | 250 (76.52) | >250 (>76.52) | >250 (>76.52) |
| RL13 | 0.24 (0.077) | 250 (80.8) | >250 (>80.8) | >250 (>80.8) |
| RL14 | 0.015 (0.0045) | 62.5 (18.69) | 250 (74.75) | 250 (74.75) |

Note: * Quantification of lanthipeptides using the eight-fold dehydrated molecules

**Table S3. Primers for construction plasmids**

| primer | Primers for PCR |
|---|---|
| **Primers of plasmids for CFPS biosynthesis** | |
| pJL1-*nisZ*-F | 5'-GGGTTTCATATGAGTACAAAAGATTTTAACTTG-3' |
| pJL1-*nisZ*-R | 5'- AATTGGATCCTTATTTGCTTACGTGAATACTAC -3' |
| pET28a-*nisB*-F | 5'-GGGTTTCATATGATAAAAAGTTCATTTAAAGC-3' |
| pET28a-*nisB*-R | 5'-GCGCGGATCCTCATTTCATGTATTCTTCCGAAAC-3' |
| pET28a-*nisC*-F | 5'-GCGCGGATCCATGAATAAAAAAAATATAAAAAG-3' |
| pET28a-*nisC*-R | 5'-ATATAAGCTTTCATTTCCTCTTCCCTCCTTTC-3' |
| pET28a-*nisP*-F | 5'-GGGTTTCATATGAAAAAAATACTAGGTTTCC-3' |
| pET28a-*nisP*-R | 5'-ATATGGATCCTCAATTTTTAGTCTTTCTTTTC-3' |
| pET28a-*nisPs*-F | 5'-AGCACCATGGATGAAAAAAATACTAGGTTTCCT-3' |
| pET28a-*nisPs*-R | 5'-<br>AGCACTCGAGTTAGTGGTGGTGGTGGTGGTGGTGTTGA<br>CTTCGTACAGAAACAGCA-3' |
| pRL-F | 5'-GCAGCCATATGAGTACAAAAGATTTTAAC-3' |
| pRL1-R | 5'- ATTCGGATCCTTAGCAGTTAACGTGGGTCG-3' |
| pRL2-R | 5'- ATTCGGATCCTTATTTGCTAATCTTGCAGTTG-3' |
| pRL3-R | 5'- ATTCGGATCCTTAGCATTTGATGCAGCTGTTG-3' |
| pRL4-R | 5'- ATTCGGATCCTTATTTGCCGGTGATTTGGCA -3' |
| pRL5-R | 5'- ATTCGGATCCTTAGCAACGAATGCAGCTGTTG-3' |
| pRL6-R | 5'- ATTCGGATCCTTACTTGCTAACGTGCACGTG-3' |
| pRL7-R | 5'- ATTCGGATCCTTAACCGGTAATGTGGATGCTG-3' |
| pRL8-R | 5'- ATTCGGATCCTTATTTGCCAATGTGGATGCTG -3' |
| pRL9-R | 5'- ATTCGGATCCTTAGCATTTAATGCAGCTGTTG -3' |
| pRL10-R | 5'- ATTCGGATCCTTATTTGCTGATGTGAACGTG-3' |
| pRL11-R | 5'- ATTCGGATCCTTATTTGCTAATGTGGCAGTTG-3' |
| pRL12-R | 5'- ATTCGGATCCTTACTTGCTCACGTGAACGTG-3' |
| pRL13-R | 5'- ATTCGGATCCTTAGTTGCCGAAGTGGCAAC-3' |
| pRL14-R | 5'- ATTCGGATCCTTATTTCTTGCCAATACCCAC-3' |
| pRL15-R | 5'- ATTCGGATCCTTAACCAACGTGCACGCTGCA-3' |
| pRL16-R | 5'- ATTCGGATCCTTAGCCAACGTGCACGCTGCA-3' |
| pRL17-R | 5'- ATTCGGATCCTTATTTGGTGTACACGGTGTC-3' |
| pRL18-R | 5'- ATTCGGATCCTTATTTGCTGATCTTGCAGCCA-3' |
| pJL1-bagelicin-R | 5'- ATTCGGATCCTTAGAAGTGGCAACCGCAGGT-3' |
| **Primers of plasmids for mlanA overexpression in *E. coli*** | |
| pYZ82-F | 5'-GCAACATATGATAAAAAGTTCATTTAAAGCTCAA-3' |
| pYZ82-R | 5'-AGCAGGTACCTCATTTCATGTATTCTTCCGAAAC -3' |
| pYZ(85-89)-F | 5'-AGCAGGATCCAATGAGTACAAAAGATTTTAACTTG-3' |
| pYZ85-R | 5'-CGATGAATTCTTACTTGCTAACGTGCACGT-3' |
| pYZ86-R | 5'-CGATGAATTCTTATTTGCCAATGTGGATGC-3' |
| pYZ87-R | 5'-AGCAGAATTCTTACTTGCTCACGTGGATCGCGCA-3' |
| pYZ89-R | 5'-AGCAGAATTCTTATTTGCTTACGTGAATCTGACA-3' |
| pYZ90-R | 5'-AGCAGAATTCTTATTTGCTTACGTGAATATCACA-3' |
| pYZ91-R | 5'-AGCAGAATTCTTATTTGCTTACGTGAATACTAC-3' |
| Sumo-F | 5'-GATATACCATGGGTCATCAC-3' |
| Sumo-R | 5'-GCTAGGATCCATATCAGCAGCGGCGCCC-3' |
| pYZ81-F | 5'-AGCAGGTACCATGAATAAAAAAAATATAAAAAGA-3' |
| pYZ81-R | 5'-AGCTCTCGAGTCATTTCCTCTTCCCTCCTTTC-3' |
| pYX125-F | 5'-CGGGGTACCATGATAAAAAGTTCATTTAAAGCTC-3' |
| pYX125-R | 5'-CCGCTCGAGTCATTTCATGTATTCTTCCGAAAC-3' |
| pYX126-F | 5'-CCGCTCGAGTGCTTAAGTCGAACAG-3' |
| pYX126-R | 5'-CCGCTCGAGTCATTTCATGTATTC-3' |
| pYX105/106-F | 5'-GTACCCTCGAGTGCTTAAGTCGAACAGAAAG-3' |

| | |
|---|---|
| pYX105/106-R | 5'-CTCGACTCGAGTCATTTCCTCTTCCCTCCTTTC -3' |
| pYX122-F | 5'-GATATGGATCCAATGAGTACAAAAGATTTTAAC-3' |
| pYX122-R | 5'-AGCTCGAATTCTTAGTTGCCGAAGTGGCAAC-3' |
| pYX123-F | 5'-GATATGGATCCAATGAGTACAAAAGATTTTAAC-3' |
| pYX123-R | 5'-AGCTCGAATTCTTATTTCTTGCCAATACCCAC-3' |
| **Primers of plasmids for nisin overexpression *in L. lactic*** | |
| pRL415-F | 5'-GTAGCTTTTTAAATATGGGTCGATCTAATATCTTGATTTTCTAGTTCCTG-3' |
| pRL415-R | 5'-TCCAAGTTAAAATCTTTTGTACTCATTTTGAGTGCCTCCTTATAATTTATT-3' |
| pRL415-VF | 5'-AATAAATTATAAGGAGGCACTCAAAATGAGTACAAAAGATTTTAACTTGGA-3' |
| pRL415-VR | 5'-CAGGAACTAGAAAATCAAGATATTAGATCGACCCATATTTAAAAAGCTAC-3' |
| pRL423-F | 5'-GTAGTATTCACGTAAGCAAATAACCAAATCAAAGGATAGTATTTTGTTAG-3' |
| pRL423-R | 5'-CTTGCATGCCTGCAGGTCGACTCTAGTCATTTCATGTATTCTTCCGAAAC-3' |
| pRL423-VF | 5'-GTTTCGGAAGAATACATGAAATGACTAGAGTCGACCTGCAGGCATGCAAG-3' |
| pRL423-VR | 5'-CTAACAAAATACTATCCTTTGATTTGGTTATTTGCTTACGTGAATACTAC-3' |

Note: For plasmid construction via restriction enzyme digestion and ligation, complementary sequences were designed using the Primer Premier 5 software. Suitable restriction sites and protective bases were introduced. For plasmid construction via the Gibson cloning method, complementary sequences were designed using the Primer Premier 5 software and were flanked by the homologous sequence. The restriction sites used for cloning are underlined.

**Table S4. Plasmids for protein purification**

| Plasmids | Replication origin | Overexpressed genes | Resistance | Reference |
|---|---|---|---|---|
| **Plasmids for CFPS biosynthesis** | | | | |
| pJL1-*nisZ* | pBR322 | PT7: N-terminal His6-tagged *nisZ* | Kan | This study |
| pET28a-*nisB* | pBR322 | PT7: N-terminal His6-tagged *nisB* | Kan | This study |
| pET28a-nisC | pBR322 | PT7: N-terminal His6-tagged *nisC* | Kan | This study |
| pET28a-*nisP* | pBR322 | PT7: N-terminal His6-tagged *nisP* | Kan | This study |
| pET28a-*nisPs* | pBR322 | PT7: N-terminal His6-tagged *nisPs* | Kan | This study |
| pRL1 | pBR322 | PT7: N-terminal His6-tagged precursor peptide gene of RL1 | Kan | This study |
| pRL2 | pBR322 | PT7: N-terminal His6-tagged precursor peptide gene of RL2 | Kan | This study |
| pRL3 | pBR322 | PT7: N-terminal His6-tagged precursor peptide gene of RL3 | Kan | This study |

| | | | | |
|---|---|---|---|---|
| pRL4 | pBR322 | PT7: N-terminal His6-tagged precursor peptide gene of RL4 | Kan | This study |
| pRL5 | pBR322 | PT7: N-terminal His6-tagged precursor peptide gene of RL5 | Kan | This study |
| pRL6 | pBR322 | PT7: N-terminal His6-tagged precursor peptide gene of RL6 | Kan | This study |
| pRL7 | pBR322 | PT7: N-terminal His6-tagged precursor peptide gene of RL7 | Kan | This study |
| pRL8 | pBR322 | PT7: N-terminal His6-tagged precursor peptide gene of RL8 | Kan | This study |
| pRL9 | pBR322 | PT7: N-terminal His6-tagged precursor peptide gene of RL9 | Kan | This study |
| pRL10 | pBR322 | PT7: N-terminal His6-tagged precursor peptide gene of RL10 | Kan | This study |
| pRL11 | pBR322 | PT7: N-terminal His6-tagged precursor peptide gene of RL11 | Kan | This study |
| pRL12 | pBR322 | PT7: N-terminal His6-tagged precursor peptide gene of RL12 | Kan | This study |
| pRL13 | pBR322 | PT7: N-terminal His6-tagged precursor peptide gene of RL13 | Kan | This study |
| pRL14 | pBR322 | PT7: N-terminal His6-tagged precursor peptide gene of RL14 | Kan | This study |
| pRL15 | pBR322 | PT7: N-terminal His6-tagged precursor peptide gene of RL15 | Kan | This study |
| pRL16 | pBR322 | PT7: N-terminal His6-tagged precursor peptide gene of RL16 | Kan | This study |
| pRL17 | pBR322 | PT7: N-terminal His6-tagged precursor peptide gene of RL17 | Kan | This study |
| pRL18 | pBR322 | PT7: N-terminal His6-tagged precursor peptide gene of RL18 | Kan | This study |
| pJL1-bagelicin | pBR322 | PT7: N-terminal His6-tagged precursor peptide gene of bagelicin | Kan | This study |
| **Plasmids for mlanA overexpression in *E. coli*** | | | | |
| pYZ82 | RSF | PT7: *nisB* | Kan | This study |
| pYZ85 | RSF | PT7: *nisB* and N-terminal His6-tagged precursor peptide gene of RL6 | Kan | This study |
| pYZ86 | RSF | PT7: *nisB* and N-terminal His6-tagged precursor peptide gene of RL8 | Kan | This study |
| pYZ87 | RSF | PT7: *nisB* and N-terminal His6-tagged precursor peptide gene of S29A | Kan | This study |
| pYZ89 | RSF | PT7: *nisB* and N-terminal His6-tagged precursor peptide gene of M5 | Kan | This study |
| pYZ90 | RSF | PT7: *nisB* and N-terminal His6-tagged precursor peptide gene of M4 | Kan | This study |
| pYZ91 | RSF | PT7: *nisB* and N-terminal His6-tagged NisZ | Kan | This study |
| pYZ92 | RSF | PT7: *nisB* and N-terminal Sumo-tagged precursor peptide gene of M4 | Kan | This study |
| pYZ93 | RSF | PT7: *nisB* and N-terminal Sumo-tagged precursor peptide gene of M5 | Kan | This study |
| pYZ95 | RSF | PT7: *nisB* and N-terminal Sumo-tagged precursor peptide gene of RL6 | Kan | This study |

| | | | | |
|---|---|---|---|---|
| pYZ96 | RSF | PT7: *nisB* and N-terminal Sumo-tagged precursor peptide gene of RL8 | Kan | This study |
| pYZ97 | RSF | PT7: *nisB* and N-terminal Sumo-tagged precursor peptide gene of S29A | Kan | This study |
| pYZ99 | RSF | PT7: *nisB* and N-terminal Sumo-tagged NisZ | Kan | This study |
| pYZ81 | P15A | PT7: *nisC* | CmR | This study |
| pYX106 | RSF | PT7: *nisB* and N-terminal His-tagged precursor peptide gene of RL6, and PT7: *nisC* | Kan | This study |
| pYX125 | P15A | PT7: *nisB* | CmR | This study |
| pYX126 | P15A | PT7: *nisC* and PT7: *nisB* | CmR | This study |
| pYX105 | RSF | PT7: *nisB* and N-terminal Sumo-tagged precursor peptide gene of S29A, and PT7: *nisC* | Kan | This study |
| pYX122 | RSF | PT7: *nisB* and N-terminal Sumo-tagged precursor peptide gene of RL13, and PT7: *nisC* | Kan | This study |
| pYX123 | RSF | PT7: *nisB* and N-terminal Sumo-tagged precursor peptide gene of RL14, and PT7: *nisC* | Kan | This study |
| **Plasmids for nisin overexpression in *L. lactis*** | | | | |
| pRL415 | pWV01 | Pnis: *nisZ* | Emr | This study |
| pRL423 | pWV01 | Pnis: *nisZ* and *nisB* | Emr | This study |

**Table S5. Gene sequences of hybrid precursor peptides.**

>RL1
ATGAGTACAAAAGATTTTAACTTGGATTTGGTATCTGTTTCGAAGAAAGATTCAGGTGC
ATCACCACGCATCACCGTGCGTAGCAAGAGCCTGTGCACCCCGGGTTGCATTACCGGT
CCGCTGCGTACCTGCTACCTGTGCTTCCCGACCCACGTTAACTGCTAA

>RL2
ATGAGTACAAAAGATTTTAACTTGGATTTGGTATCTGTTTCGAAGAAAGATTCAGGTGC
ATCACCACGCATCACCTGGAAGAGCGAGAGCCTGTGCACCCCGGGTTGCGTGACCGG
CGTTCTGCAGACCTGCTTCCTGCAAACCATCACCTGCAACTGCAAGATTAGCAAATAA

>RL3
ATGAGTACAAAAGATTTTAACTTGGATTTGGTATCTGTTTCGAAGAAAGATTCAGGTGC
ATCACCACGCGTGACCAGCAAGAGCCTGTGCACCCCGGGTTGCATCACCGGCATTCTG
ATGTGCCTGACCCAGAACAGCTGCGTTAGCTGCAACAGCTGCATCAAATGCTAA

>RL4
ATGAGTACAAAAGATTTTAACTTGGATTTGGTATCTGTTTCGAAGAAAGATTCAGGTGC
ATCACCACGCATCACCAGCAAAAGCCTGTGCACCCCGGGTTGCGTGACCGGCATTCTG
ATGACCTGCCCGGTTCAGACCGCGACCTGCGGTTGCCAAATCACCGGCAAATAA

>RL5
ATGAGTACAAAAGATTTTAACTTGGATTTGGTATCTGTTTCGAAGAAAGATTCAGGTGC
ATCACCACGCATCACCAGCAAGAGCCTGTGCACCCCGGGTTGCATCACCGGCATTCTG
ATGTGCCTGACCCAGAACAGCTGCGTGAGCTGCAACAGCTGCATTCGTTGCTAA

>RL6
ATGAGTACAAAAGATTTTAACTTGGATTTGGTATCTGTTTCGAAGAAAGATTCAGGTGC
ATCACCACGCATCACCAGCGTGAGCCTGTGCACCCCGGGTTGCAAGACCGGTGCGCTG
ATGGGTTGCAACATGAAAACCGCGAGCTGCGGCTGCCACGTGCACGTTAGCAAGTAA

>RL7
ATGAGTACAAAAGATTTTAACTTGGATTTGGTATCTGTTTCGAAGAAAGATTCAGGTGC
ATCACCACGCATCACCAGCGTGAGCCTGTGCACCCCGGGTTGCGTGACCGGCGTTCTG

ATGTGCCCGGGTAACACCATTAGCTGCAACGGCCACTGCAGCATCCACATTACCGGTTA
A
>RL8
ATGAGTACAAAAGATTTTAACTTGGATTTGGTATCTGTTTCGAAGAAAGATTCAGGTGC
ATCACCACGCGTGACCAGCAAGAGCCTGTGCACCCCGGGTTGCAAAACCGGCATCCT
GCAGACCTGCGCGATTAAGAGCGCGACCTGCGGTTGCAGCATCCACATTGGCAAATAA
>RL9
ATGAGTACAAAAGATTTTAACTTGGATTTGGTATCTGTTTCGAAGAAAGATTCAGGTGC
ATCACCACGCGTGACCAGCAAGAGCCTGTGCACCCCGGGTTGCATCACCGGCGTGCTG
ATGTGCCTGACCCAGAACAGCTGCGTTAGCTGCAACAGCTGCATTAAATGCTAA
>RL10
ATGAGTACAAAAGATTTTAACTTGGATTTGGTATCTGTTTCGAAGAAAGATTCAGGTGC
ATCACCACGCATCACCGTGAAGATTACCAGCTACAGCCTGTGCACCCCGGGTTGCAAG
ACCGGTGCGCTGATGGGTTGCACCATGAAAACCGCGAGCTGCGGCTGCCACGTTCACA
TCAGCAAATAA
>RL11
ATGAGTACAAAAGATTTTAACTTGGATTTGGTATCTGTTTCGAAGAAAGATTCAGGTGC
ATCACCACGCATCACCTGGAAGAGCGAGAGCCTGTGCACCCCGGGTTGCATTACCGGC
GTGCTGCAGACCTGCTTCCTGCAAACCATCACCTGCAACTGCCACATTAGCAAATAA
>RL12
ATGAGTACAAAAGATTTTAACTTGGATTTGGTATCTGTTTCGAAGAAAGATTCAGGTGC
ATCACCACGCATCACCAGCTACAGCCTGTGCACCCCGGGTTGCATTACCGGCGTTCTGA
TGGGTTGCCACATCCAGAGCATTGGCTGCAACGTGCACGTTCACGTGAGCAAGTAA
>RL13
ATGAGTACAAAAGATTTTAACTTGGATTTGGTATCTGTTTCGAAGAAAGATTCAGGTGC
ATCACCACGCATCACCAGCAAGAGCCTGTGCACCCCGGGTTGCAAAACCGGCGCGCT
GATGACCTGCCCGATTAAGACCGCGACCTGCGGTTGCCACTTCGGCAACTAA
>RL14
ATGAGTACAAAAGATTTTAACTTGGATTTGGTATCTGTTTCGAAGAAAGATTCAGGTGC
ATCACCACGCATCACCAGCAAGAGCCTGTGCACCCCGGGTTGCGTGACCGGCGTTCTG
ATGGGTTGCGCGCTGAAAACCATCACCTGCAACTGCAGCGTGGGTATTGGCAAGAAAT
AA
>RL15
ATGAGTACAAAAGATTTTAACTTGGATTTGGTATCTGTTTCGAAGAAAGATTCAGGTGC
ATCACCACGCATCACCAGCAAGAGCCTGTGCACCCCGGGTTGCGTTACCGGTCTGCTG
ATGGGTTGCGCGGGTAGCAGCGCGACCTGCAACTGCAGCGTGCACGTTGGTTAA
>RL16
ATGAGTACAAAAGATTTTAACTTGGATTTGGTATCTGTTTCGAAGAAAGATTCAGGTGC
ATCACCACGCATCACCAGCAAGAGCCTGTGCACCCCGGGTTGCGTGACCGGCGTTCTG
ATGGGTTGCAACAACAAAACCGCGACCTGCAACTGCAGCGTGCACGTTGGCTAA
>RL17
ATGAGTACAAAAGATTTTAACTTGGATTTGGTATCTGTTTCGAAGAAAGATTCAGGTGC
ATCACCACGCATCACCCAGTTCAAGAGCATTAGCCTGTGCACCCCGGGTTGCCCGACC
GGTATCCTGATGGGTTGCCATAAGTGCCCGAGCGGTAGCGACACCGTGTACACCAAAT
AA
>RL18
ATGAGTACAAAAGATTTTAACTTGGATTTGGTATCTGTTTCGAAGAAAGATTCAGGTGC
ATCACCACGCATCACCAGCCCGCAGATTACCAGCGTGAGCCTGTGCACCCCGGGTTGC
CAGACCGGCTTCCTGGCGTGCTTTAGCCAAGCGTGCAACCCGACCGGTGGCTGCAAG
ATCAGCAAATAA
>NisZ
ATGAGTACAAAAGATTTTAACTTGGATTTGGTATCTGTTTCGAAGAAAGATTCAGGTGC
ATCACCACGCAGTATTTCGCTATGTACACCCGGTTGTAAAACAGGAGCTCTGATGGGTT
GTAACATGAAAACAGCAACTTGTAATTGTAGTATTCACGTAAGCAAATAA
> Bagelicin
ATGAGTACAAAAGATTTTAACTTGGATTTGGTATCTGTTTCGAAGAAAGATTCAGGTGC
ATCACCACGCGTGACCAGCATCAGCCTGTGCACCCCGGGTTGCAAGACCGGCATCCTG

ATGACCTGCGCGATTAAAACCGCGACCTGCGGTTGCCACTTCTAA
>M4
ATGAGTACAAAAGATTTTAACTTGGATTTGGTATCTGTTTCGAAGAAAGATTCAGGTGC
ATCACCACGCATTACAAGTAAGTCGCTATGTACAACCGGTTGTCTGACAGGAGCGCTG
ATGGGTTGTAACATGAAAACAGCGACTTGTAATTGTGATATTCACGTAAGCAAATAA
>M5
ATGAGTACAAAAGATTTTAACTTGGATTTGGTATCTGTTTCGAAGAAAGATTCAGGTGC
ATCACCACGCATTACAAGTCGGTCGCTATGTACACCCGGTTGTTGGACAGGACCTCTGA
TGGGTTGTAACATGAAAACAAAGACTTGTAATTGTCAGATTCACGTAAGCAAATAA
>S29A
ATGAGTACAAAAGATTTTAACTTGGATTTGGTATCTGTTTCGAAGAAAGATTCAGGTGC
ATCACCACGCATCACCAGCATTAGCCTGTGCACCCCGGGTTGCAAGACCGGTGCGCTG
ATGGGTTGCAACATGAAAACCGCGACCTGCCACTGCGCGATCCACGTGAGCAAGTAA

Note: leader peptide sequence is gray marked, core peptide sequence is underlined.

**Reference**


1. G. Y. Tan, K. H. Deng, X. H. Liu, H. Tao, Y. Y. Chang, J. Chen, K. Chen, Z. Sheng, Z. X. Deng, T. G. Liu, Heterologous Biosynthesis of Spinosad: An Omics-Guided Large, Polyketide Synthase Gene Cluster Reconstitution in *Streptomyces*. *Acs Synth Biol* **6**, 995-1005 (2017).
2. B. Soufi, F. Gnad, P. R. Jensen, D. Petranovic, M. Mann, I. Mijakovic, B. Macek, The Ser/Thr/Tyr phosphoproteome of *Lactococcus lactis* IL1403 reveals multiply phosphorylated proteins. *Proteomics* **8**, 3486-3493 (2008).
3. D. Field, M. Begley, P. M. O'Connor, K. M. Daly, F. Hugenholtz, P. D. Cotter, C. Hill, R. P. Ross, Bioengineered nisin A derivatives with enhanced activity against both Gram positive and Gram negative pathogens. *Plos One* **7**, e46884 (2012).
4. L. Zhou, A. J. van Heel, M. Montalban-Lopez, O. P. Kuipers, Potentiating the activity of nisin against *Escherichia coli. Front Cell Dev Biol* **4**, 7 (2016).
5. Q. Li, M. Montalban-Lopez, O. P. Kuipers, Increasing the antimicrobial activity of nisin-based lantibiotics against Gram-negative pathogens. *Appl Environ Microbiol* **84**, (2018).
